# Diverse priming outcomes under conditions of very rare precursor B cells

**DOI:** 10.1101/2024.11.21.624746

**Authors:** Patrick J. Madden, Ester Marina-Zárate, Kristen A. Rodrigues, Jon M. Steichen, Monolina Shil, Kaiyuan Ni, Katarzyna Kaczmarek Michaels, Laura Maiorino, Amit A. Upadhyay, Swati Saha, Arpan Pradhan, Oleksandr Kalyuzhiny, Alessia Liguori, Paul G. Lopez, Ivy Phung, Nicole Phelps, Erik Georgeson, Nushin Alavi, Michael Kubitz, Danny Lu, Saman Eskandarzadeh, Amanda Metz, Oscar L. Rodriguez, Kaitlyn Shields, Steven Schultze, Melissa L. Smith, Brandon S. Healy, Deuk Lim, Vanessa R. Lewis, Elana Ben-Akiva, William Pinney, Justin Gregory, Shuhao Xiao, Diane G. Carnathan, Sudhir Pai Kasturi, Corey T. Watson, Steven E. Bosinger, Guido Silvestri, William R. Schief, Darrell J Irvine, Shane Crotty

**Affiliations:** Center for Vaccine Innovation, La Jolla Institute for Immunology, La Jolla, CA, USA; Consortium for HIV/AIDS Vaccine Development (CHAVD), The Scripps Research Institute, La Jolla, CA, USA; Koch Institute for Integrative Cancer Research, Massachusetts Institute of Technology, Cambridge, MA, USA; Department of Immunology and Microbiology, The Scripps Research Institute, La Jolla, CA, USA; Emory National Primate Research Center and Emory Vaccine Center, Emory University School of Medicine, Atlanta, GA, USA; Department of Pathology and Laboratory Medicine, Emory School of Medicine, Atlanta, GA, USA; Department of Biochemistry and Molecular Genetics, University of Louisville School of Medicine, Louisville, KY, USA; IAVI Neutralizing Antibody Center, The Scripps Research Institute, La Jolla, CA, USA; Moderna, Inc., Cambridge, MA, USA; Department of Biological Engineering, Massachusetts Institute of Technology, Cambridge MA USA; Howard Hughes Medical Institute, Chevy Chase, MD, USA; Department of Medicine, Division of Infectious Diseases and Global Public Health, University of California, San Diego (UCSD), La Jolla, CA, USA

## Abstract

Rare B cells can have special pathogen-recognition features giving them the potential to make outsized contributions to protective immunity. However, rare naive B cells infrequently participate in immune responses. We investigated how germline-targeting vaccine antigen delivery and adjuvant selection affect priming of exceptionally rare BG18-like HIV broadly neutralizing antibody-precursor B cells (~1 in 50 million) in non-human primates. Only escalating dose (ED) priming immunization using the saponin adjuvant SMNP elicited detectable BG18-like cells in germinal centers (GCs). All groups had strong GC responses, but only ED+SMNP and bolus+SMNP induced BG18-like memory B cells in >50% of animals. One group had vaccine-specific GC responses equivalent to ED+SMNP, but BG18-like memory B cells were rarely detected. Following homologous boosting, BG18-like memory B cells were more frequent in a bolus priming group, but had lower somatic hypermutation and affinities. This outcome was inversely associated with post-prime antibody titers, suggesting antibody feedback can significantly influence rare precursor B cell responses.

## INTRODUCTION

The aim of many vaccines against viruses is to induce neutralizing antibodies that protect against infection^1–3^. The B cell precursors that give rise to these types of neutralizing responses can be extremely rare in the naïve B cell repertoire and infrequently participate in immune responses^4–12^. Germline-targeting vaccine design is one approach to prime these rare B cell specificities by designing immunogens that have affinity for the precursors of choice^13–16^. HIV broadly neutralizing antibody (bnAb) precursors have been a main focus of germline-targeting immunogen design^5,8,12,17–28^. This approach relies on the concept that immunogens that can be recognized with high affinity by rare bnAb-precursor B cells can overcome their inherent rarity and allow for sufficient B cell activation and recruitment into germinal center (GC) reactions^12,19,29^. Using adoptive transfer experiments in mice to model human precursor frequencies, rare bnAb-precursor B cells can be successfully primed if the immunogen has sufficient affinity for the target naïve B cell receptors (BCRs)^4^. These outcomes have been tested and validated in a first-in-humans germline-targeting vaccine clinical trial wherein rare VRC01-class bnAb precursors were primed in 97% of vaccine recipients^26^.

It was recently shown that a novel HIV germline-targeting immunogen, N322-GT5, designed to target and prime rare precursors similar to the HIV bnAb BG18, successfully primed BG18-class responses in 8 out of 8 rhesus macaques (RMs) ^12,19,25^. In humans, N332-GT5 reactive BG18-class precursors were found at a frequency of 1 in 53 million in the naïve B cell repertoire^12^, and are estimated to be 8-fold rarer in RM^25^. In addition to being an important advance in HIV candidate vaccine development in the germline-targeting vaccine field, the success of N332-GT5 in NHPs provides a new opportunity for a model system to study rare B cell priming (defined as precursor frequencies < 1 in 1 million naive B cells) in outbred, non-transgenic animals. It is particularly advantageous to be able to do so in a primate species, given the much more similar immunoglobulin gene repertoire features and heavy chain complementarity determining region 3 (H-CDR3) lengths between non-human primates and humans compared to rodents^30,31^.

It is yet to be elucidated how adjuvants and antigen delivery strategies affect the recruitment of rare B cells into GCs, despite much activity in adjuvant development and new vaccine technologies^32–36^. To build on the success of germline targeting immunogen design and explore the immunology of rare B cell recruitment, here we investigated the impact of various delivery strategies and adjuvants on the priming of BG18-class precursor B cells in RMs. The results reveal key features about successful versus failed priming and maturation of rare naive B cells in different contexts.

## RESULTS

### Adjuvant/delivery strategies and vaccine study design

To assess whether different immunization strategies affect recruitment of rare bnAb-precursor B cells in the context of a germline-targeting priming immunization, 36 RMs were split into six groups (n = 6/group) and immunized with the recombinant HIV Env-based germline-targeting immunogen N332-GT5 (**Fig. 1**). ED immunization is a slow antigen delivery immunization technique^37,38^. Saponin and monophosphoryl lipid A (MPLA)-containing nanoparticle (SMNP) is a potent adjuvant^25,39^ being developed for clinical use^40^. ED immunization adjuvanted with other immune stimulating complexes (ISCOMs) has been previously shown to recruit a more diverse set of naïve B cells to participate in GCs^38^, including rare neutralizing antibody precursor B cells, when compared to traditional bolus immunization^37^. ED immunization adjuvanted with SMNP (ED + SMNP) successfully primed BG18-class B cells in NHPs^25^.

**Figure 1:**
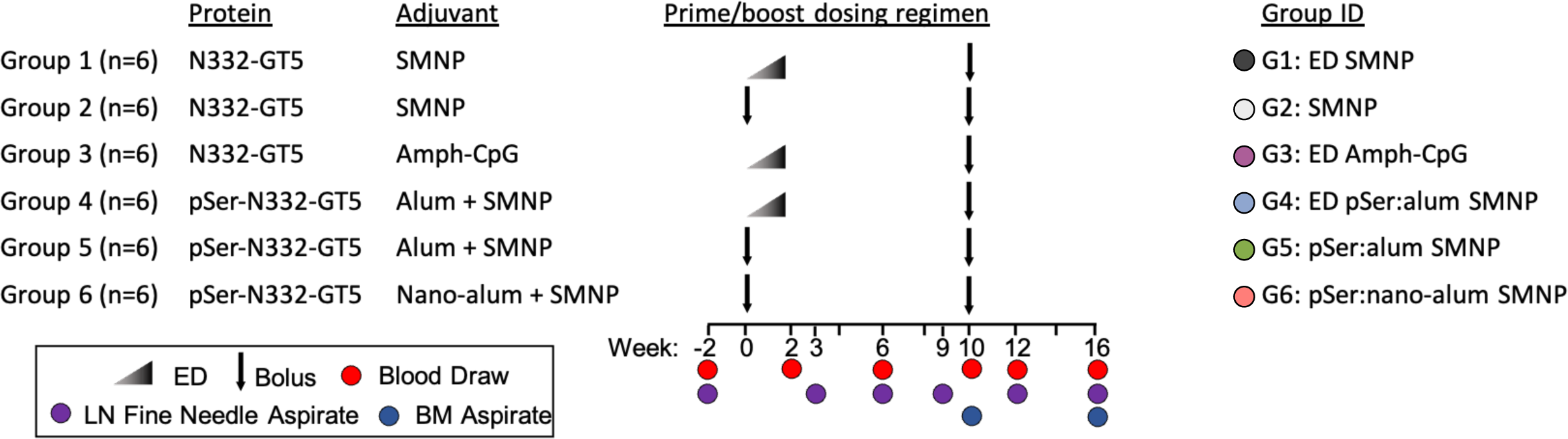
Study schematic. Study schematic showing seven immunization groups and sampling types and timepoints (n =6 for each)

We hypothesized that the extended-antigen availability afforded by ED immunization may lead to better recruitment and priming of BG18-class B cells^25,41^. An additional method to extend antigen availability is via adsorption of phosphoserine-tagged immunogens on aluminum hydroxide (alum) (pSer:alum)^42^. Tethering immunogens to alum particles in this way slows antigen clearance while simultaneously increasing the potential avidity of the immunogen^42–44^. Previously, we have shown that pSer:alum delivery of a native-like HIV Env trimer in RMs enhanced responses compared to unmodified antigen with alum, and adjuvanting the pSer:alum complexes with SMNP led to even better responses^45^. The bnAb BG18 targets the V3/glycan site on HIV Env, which includes the N322 glycan. By orienting the N332-GT5 Env trimer on alum, it may promote responses to the V3/glycan bnAb epitopes and reduce off-target Env base-binding responses^42,43,45^. We thus synthesized N332-GT5 trimers with an optimized C-terminal Cys linker^43^, and coupled a peptide containing 4 phosphoserines to the base of each protomer to enable pSer-mediated anchoring to alum (**Supp. Fig. 1A**). pSer-modified N332-GT5 showed stable binding to alum in the presence of serum and exhibited high binding of HIV bnAbs (particularly inferred-germline BG18) and low binding of trimer base-specific Abs, as expected (**Supp. Fig. 1B-D**).

Alum is a suspension of aluminum hydroxide nanocrystals that aggregate to form particles of varying sizes^42,46,47^. Stable dispersions of sub-micron sized aluminum hydroxide particles, termed nano-alum, have been shown to elicit stronger antibody responses against multiple vaccines when compared to normal alum^48^. To test the potential of nano-alum in the setting of germline targeting, we generated nano-alum by dispersing Alhydrogel in the presence of a poly(ethylene glycol) (PEG) phospholipid. The PEG-lipid bound to the alum through its phosphate group, allowing individual needle-like aluminum hydroxide crystals to be stably dispersed in buffer (**Supp. Fig. 2A**). Nano-alum had a particle size under 100 nm diam. as assessed by dynamic light scattering and individual rod-like nanocrystals observed in TEM, which were smaller than the microscale aggregates of alum particles in unmodified Alhydrogel (**Supp. Fig. 2B-C**). While unmodified alum slowly sediments in water, nano-alum formed a stable opalescent solution (**Supp. Fig. 2D**). When mixed with pSer-trimer, many Env trimers could be observed decorating the length of individual nano-alum crystals by TEM (**Supp. Fig. 2E**). In preliminary immunization studies in mice, nano-alum loaded with pSer-modified Env trimers elicited substantial increases in antigen-specific GC B cells and serum IgG titers compared to traditional alum loaded with pSer-trimer (**Supp. Fig. 2F-H**).

Toll-like receptor (TLR) agonists are also being widely studied as vaccine adjuvants^33,34,36,39^, and this study also evaluated multiple TLR agonist formulations for their potential to recruit rare B cell precursors. The MPLA in SMNP is a TLR-4 agonist. CpG is a TLR9 agonist that can be further enhanced by the addition of an albumin-binding lipid tail (Amph-CpG, **Supp. Fig. 3**)^49^. Amph-CpG takes advantage of the fact that large proteins, such as albumin, cannot efficiently pass from tissue to blood and therefore must enter lymphatics^49^. This leads to increased uptake of the amph-CpG in draining lymph nodes (LNs) by bypassing systemic distribution in blood^49^. Amph-CpG has been shown to be a potent adjuvant in RMs immunized with a SARS-CoV-2 protein vaccine^50^.

The six study groups comprised different combinations of three delivery strategies (bolus, ED, and pSer) and four adjuvants (SMNP, alum, nano-alum, Amph-CpG) and are detailed in **Fig. 1**. Each of these adjuvants have distinct properties and have been highly successful in different contexts, but they have not been tested in the context of rare precursor B cell recruitment. All immunizations were administered subcutaneously in each thigh (50μg protein/side, 100μg total) at weeks 0 and 10 (**Fig. 1A**). The week 10 boosting immunizations were all bolus, using the same immunogen plus adjuvant combination and dose used at the priming timepoint by group. To date, only ED SMNP delivery of N332-GT5 has been tested and shown to prime BG18 type I precursor cells. A primary objective of this study was to thus determine the immunology of rare bnAb-precursor B cell priming under a range of conditions. Two secondary objectives were also explored to determine whether the pSer:alum based immunizations were superior to traditional bolus immunization, and to assess how pSer:alum behaved when delivered in an ED immunization with different adjuvants. Statistical tests presented here were designed to reflect these specific objectives and are detailed in the methods.

### Escalating dose immunizations and SMNP prime larger GC responses

GC responses were measured by flow cytometry from lymph node fine needle aspirates (LN FNAs) at multiple timepoints throughout the study. ED immunizations led to significantly larger total GC B cell (B_GC,_ CD38^−^CD71^+^) and GC-T_FH_ (PD-1^Hi^CXCR5^+^) responses post-prime (weeks 3-9) when compared to bolus groups using the same adjuvant combinations (G1 vs G2, and G4 vs G5, **Fig. 2A-F**). G3 had lower B_GC_ and GC-T_FH_ responses throughout the study compared to G1 and G4, indicating that SMNP was more effective in driving GC responses than Amph-CpG in the setting of escalating-dose immunization (**Fig. 2B, E**).

**Figure 2:**
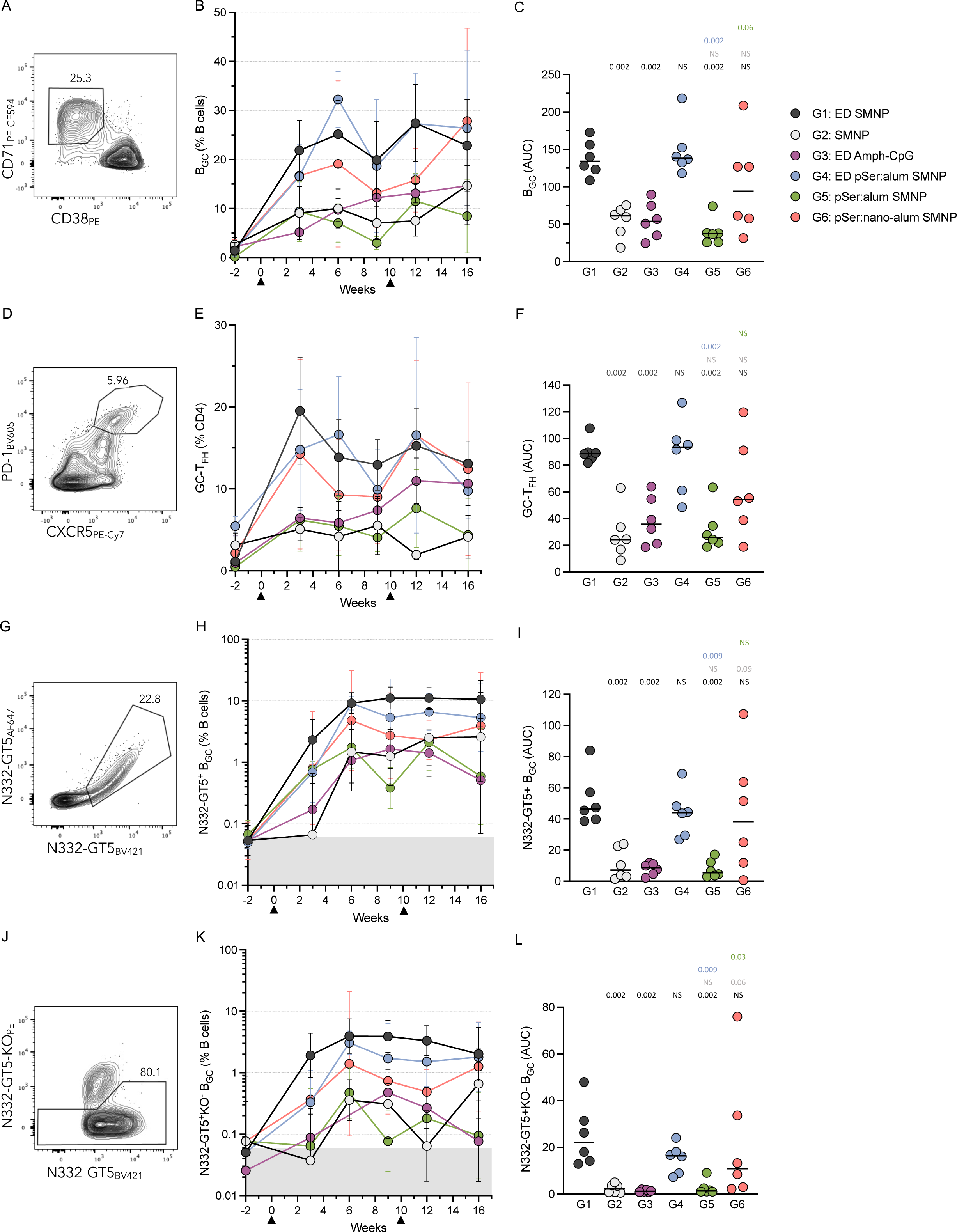
Escalating dose immunization and SMNP prime larger GC responses. **A)** Flow cytometry gating of B_GC_ cells (CD38^−^CD71^+^). **B)** B_GC_ cell frequency (CD38^−^CD71^+^) as a percentage of total B cells (CD20^+^). **C)** Area under the curve (AUC) of B_GC_ cell frequency as a percentage of total B cells post-priming (weeks 3-9) immunization. **D)** Flow cytometry gating of GC-T_FH_ cells (PD-1^Hi^CXCR5^+^). **E)** GC-T_FH_ cell frequency as a percentage of total CD4^+^ cells. **F)** AUC of GC-T_FH_ cell frequency as a percentage of total CD4^+^ cells post-priming (week 3-9) immunization. **G)** Flow cytometry gating of N332-GT5 antigen specific B_GC_ cells (N332-GT5-AF647^+^N332-GT5-BV421^+^; N332-GT5^++^). **H)** Antigen-specific B_GC_ cell frequency (N332-GT5-AF647^+^N332-GT5-BV421^+^) as a percentage of total B cells (CD20^+^). **I)** Area under the curve (AUC) of antigen-specific B_GC_ cell frequency as a percentage of total B cells post-priming (weeks 3-9) immunization. **J)** Flow cytometry gating of N332-GT5 epitope specific B_GC_ cells (N332-GT5-AF647^+^N332-GT5-BV421^+^N332-GT5KO-PE^−^). **K)** Epitope-specific B_GC_ cell frequency (N332-GT5-AF647^+^N332-GT5-BV421^+^N332-GT5KO-PE^−^) as a percentage of total B cells (CD20^+^). **I)** Area under the curve (AUC) of epitope-specific B_GC_ cell frequency as a percentage of total B cells post-priming (weeks 3-9) immunization. Triangles in longitudinal graphs represent time of immunizations. Mean and SEM or geometric mean and geometric SD are plotted depending on the scale in all longitudinal figures and median was plotted in all per animal figures. Statistical significance was tested using unpaired two-tailed Mann-Whitney tests with the p-values listed on each graph representative of the tests carried out, NS listed when p-value was >0.1. Gray regions (H, K) represent B_GC_ LOD determined using pre-immunization samples.

B_GC_ cells were also stained with fluorophore-labeled N332-GT5 probes to detect antigen-specific (two colors of N332-GT5 probes; N332-GT5^++^) and epitope-specific (N332-GT5^++^N332-GT5KO^−^) B_GC_ cells (**Fig. 2G-L**). All groups had detectable antigen- and epitope-specific B_GC_ cells starting at week 3, but the kinetics of GC priming differed (**Fig. 2G-L**). G1 and G4 had the largest post-prime antigen- and epitope-specific B_GC_ frequencies. At week 3, bolus SMNP vaccination (G2) elicited a frequency of antigen-specific B_GC_ cells barely above the limit of detection (LOD, baseline determined pre-immunization), followed by a sharp rise at week 6, which was mirrored by the expansion of epitope-specific B_GC_ cells (**Fig. 2H, K**). Interestingly, all of the groups employing SMNP in an extended-delivery regimen (either ED or pSer) expanded antigen-specific B_GC_ cells 10-fold or more by week 3 (**Fig. 2H**). This was most pronounced in G1, where by week 3 both antigen- and epitope-specific B_GC_ cells were rapidly increased 30-40-fold above baseline. Despite the rise at week 6, G2 still had a ~10-fold lower frequency of epitope-specific B_GC_ cells at weeks 6 and 9 compared to G1 (**Fig. 2K**, p = 0.002 week 6 and 0.002 week 9, Mann-Whitney). When looking at total post-prime responses, G1 had significantly higher AUC values for both antigen- and epitope-specific B_GC_ cells compared to all groups except G4 and G6 (**Fig. 2I, L**). These data suggest that ED SMNP generated robust GCs faster than other groups and recruited epitope-specific cells to GCs to a greater extent than all but G4 and G6.

We also examined the antigen- and epitope-specific non-B_GC_ cells at weeks 3 and 9 to determine if certain groups drove a more extrafollicular or memory response in lymph nodes. The hierarchy of the non-B_GC_ cell responses for the animal groups mirrored that seen for the B_GC_ for both antigen- and epitope-specific cells (**Supp. Fig 4B-C**).

Unexpectedly, the slow-delivery behavior of pSer:alum combined with SMNP immunization did not enhance total GC or antigen-specific B_GC_ cell responses over SMNP alone in either bolus or ED administration. However, pSer-conjugated trimer delivered on nano-alum led to significantly more epitope-specific B_GC_ cells out to week 10 compared to traditional alum (G6 vs G5) and was the only bolus-administered formulation to increase GC responses approaching ED dosing (**Fig. 2I, L**), suggesting that pSer:nano-alum has a unique mechanism of action.

Overall, these data from 36 animals over a 10-week time course indicate that the two ED SMNP containing immunizations primed the most robust total and antigen-specific GC responses, with notably accelerated B_GC_ expansion compared to bolus vaccination. Enhanced frequencies of epitope-specific B_GC_ suggested that ED immunization with SMNP can better recruit diverse B cells after priming compared to bolus or non-SMNP adjuvanted immunizations.

### Escalating dose adjuvanted with SMNP leads to a larger and more diverse BCR response

Epitope-specific B_GC_ cells were FACS sorted and single cell BCR sequencing was completed at week 3 post-immunization to assess the ability of different delivery and adjuvant combinations to recruit and prime rare BG18 type I precursor cells. A total of 26,207 epitope-specific paired BCRs from week 3 LN FNAs were sequenced across all groups. BG18 type I BCRs are defined based on immunoglobulin heavy chain (HC) sequence features and H-CDR3 length (**Supp. Fig. 5A**). BG18 type I BCRs were detected in 3/6 animals from G1 but no animals in any other group (**Fig. 3A, Supp.Table 1**). The results from G1 were consistent with the original N332-GT5 RM study, in which 5/8 ED SMNP animals had BG18 type I B_GC_ cells at week 3 (ref. ^25^). G4 and G6 had similar total post-prime antigen- and epitope-specific B_GC_ cells compared to G1 (**Fig. 2H, K**), thus it was surprising that no BG18 type I BCRs were found in these groups. If BG18 type I precursor priming was equivalent in all groups, BG18 type I BCRs were expected in all groups except the non-SMNP group (G3) based on total sequencing per group (**Fig. 3A**). These data indicate that only ED SMNP immunization possessed properties necessary for successful recruitment and expansion of very rare BG18 type I precursor cells above a limit of detection of ~1 in 3,000 B_GC_ by 3-weeks post-vaccination.

**Figure 3:**
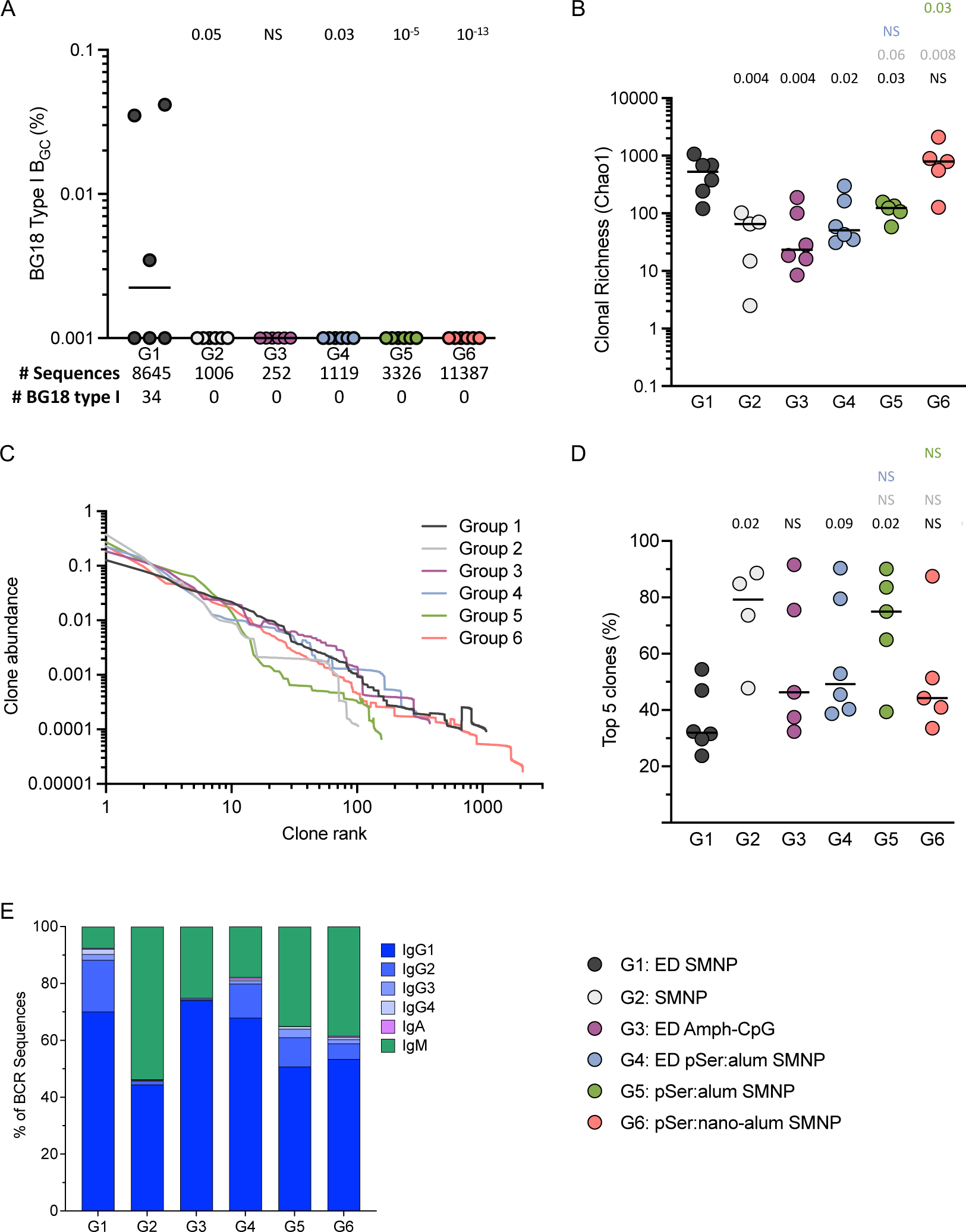
Escalating dose adjuvanted with SMNP leads to a larger, more diverse, composition of BCRs 3-weeks post-prime. **A)** Frequency of BG18 type I B_GC_ cells from week 3 among total B cells, plotted per animal. Numbers below indicate total number of paired BCR sequences recovered from each group and the total number of BG18 type I BCRs recovered. All cells are from week LN FNAs. **B)** Clonal richness of the BCR sequences recovered from epitope specific B_GC_ cells plotted as the Chao1 index on a per animal basis. Chao was calculated for all animals that had at least 1 sequence recovered. **C)** Clonal abundance curves for each group. Clonal abundance was calculated per animal then the mean abundance for each clone rank was plotted for each group. Animals with fewer than 5 total clones were excluded from abundance calculations. **D)** Cumulative abundance of the top 5 most abundant clones plotted per animal. Animals with fewer than 5 total clones were excluded from abundance calculations. **E)** Percentage of each Ig isotype making up the total BCR sequences isolated from each group. The lines in all per animal graphs represent medians. Statistical significance was tested using unpaired two-tailed Mann-Whitney tests with the p-values listed on each graph representative of the tests carried out except for A) where multiple Fisher’s exact tests were used to compare each group directly to G1, NS listed when p-value was >0.1.

As an additional measure of rare B cells, we searched for two other populations of cells; cells similar to BG18 type I cells but with shorter H-CDR3 length, and potential BG18 type III cells. The original definition used for BG18 type I cells in RMs includes H-CDR3s of at least 22 amino acids with additional specific H-CDR3 sequence features^12^, but we observed BCRs that otherwise meet the criteria for being BG18 type I but have H-CDR3s of 20 or 21 AAs (“BG18_20-21AA_” **Supp. Fig. 5A-B**). When quantified, BG18_20-21AA_ cells were found in the same three animals from G1 where the traditional BG18 type I cells were found but at a slightly lower frequency (**Supp. Fig. 5B**). We also observed a single BG18_20-21AA_ BCR in one animal each from G2 and G3 despite no traditional BG18s being found. BG18 type III BCRs, defined as those having H-CDR3 lengths ≥20 AAs and a binding angle of approach and footprint similar to BG18 type I but lacking other sequence features of BG18 type I or type II, were previously identified from RMs immunized with N332-GT5 using B cell sorting, Fab production, and cryoEM studies^25^. We searched the BCRs for sequences that used the same IGHD gene and contained the important contact residues from the cryoEM structures as potential type III cells. These cells were found at similar frequencies to the traditional and BG18_20-21AA_ responses but were identified in two additional animals (**Supp. Fig. 5C).** These results further corroborate that rare potential bnAb-precursor B cells are recruited and expanded after ED SMNP immunization better than other immunization strategies.

To further understand the composition of the B_GC_ response at the population level, clonal richness (Chao1) and diversity (Simpson index) were calculated of the epitope-specific B_GC_. ED SMNP elicited a larger and more diverse population of B_GC_ cells compared to all others groups except G6 (**Fig. 3B, Supp.Fig. 5D**). There were no significant differences between pSer:alum groups G2, G4, and G5 in either clonal richness or diversity, indicating that pSer:alum delivery, by bolus or ED, offered no clear advantage over bolus immunization for increasing on-target B cell diversity (**Fig. 3B, Supp.Fig. 5D**). In contrast, G6 (nano-alum) had clonal richness comparable to G1 and significantly increased clonal richness compared to G5 (**Fig. 3B**), indicating that pSer:nano-alum changed the overall B cell composition of the GC responses compared to pSer:alum.

Clonal abundance curves were plotted for each group (**Fig. 3C**), and the cumulative abundance of the top 5 clones was calculated for each animal (**Fig. 3D**). For G1, the top 5 clones accounted for ~30% of the total B_GC_ response (median per group), while for G2 the median was significantly higher at ~80% (**Fig. 3D**). Together, these parameters show that ED SMNP recruited large numbers of clones with moderated immunodominance, whereas bolus SMNP immunization recruited fewer total B cell clones and a small number of those clones dominated the GC response.

Ig isotype was determined for each sequenced BCR (**Fig. 3E**). All ED groups had a higher frequency of class-switched isotypes (IgG1-4, IgA) than bolus immunization. For G2, over 50% of the epitope specific B_GC_ cells at week 3 were IgM. This was in contrast to 90% of the epitope-specific B_GC_ cells being IgG class-switched in G1 (**Fig. 3E**). When interpreted with the GC kinetics data (**Fig. 2H, K**), the class switch frequency differences suggest that the GC responses were delayed after bolus immunization compared to ED SMNP (**Fig. 3E**). Despite the increased size and diversity of the GC response for G6 compared to G5, both groups had similar isotype distributions, indicating that GC kinetics may be similar in both groups, and the differential mechanism of nano-alum may lie more in the total number of different B cells recruited to GCs.

Overall, the antigen-specific BCR sequencing data showed that in GCs 3-weeks post-prime, only ED SMNP recruited detectable BG18 type I cells, which correlated with a distinctly clonally rich, diverse, and rapid B_GC_ cell response compared to other groups. Regarding kinetics of the GCs, antigen- and epitope-specific B_GC_ frequencies showed that many groups still had low vaccine-specific GC responses at the 3-week time point, which subsequently peaked at 6 weeks post-prime, or even 10 weeks post-prime (**Fig 2H, K**). It therefore remained possible that the clonal composition of those more slowly amplifying B cell responses may evolve after week 3.

### Vaccine-specific T cell responses

Germinal center responses require CD4 T cell help, specifically T_FH_ cells^51^. N332-GT5 specific T cell responses were measured by activation-induced marker (AIM) and intracellular cytokine staining (ICS) assays at 2-weeks post-immunization for multiparametric enumeration and functional analysis of the CD4 T cell response (**Fig. 4A**). N332-GT5–specific (AIM^+^, CD40L^+^OX40^+^) CD4^+^ T cell responses were detected in all six groups above the pre-immunization levels (**Fig. 4B**). The three pSer:alum groups had the lowest T cell responses (G4-G6, **Fig. 4B**). Similar trends between groups were seen with other AIM parameters, and by ICS for IFNγ, granzyme B, IL2, and TNF (**Supp.Fig. 6B-F**). Antigen-specific circulating T_FH_ (cT_FH_) cells were measured (**Fig. 4C**). G1 had a significantly higher frequency of antigen-specific (AIM^+^CD40L^+^OX40^+^) cT_FH_ cells compared to all other groups (**Fig. 4D, Supp.Fig. 6G**). The remaining groups had similar frequencies of cT_FH_ cells, despite clear differences in GC-T_FH_ frequencies between groups. For G4 and G6, both had ~3-fold higher frequencies of GC-T_FH_ compared to other groups but similar levels of cT_FH_ (**Fig. 4D, 2E-F,** week 3).

**Figure 4:**
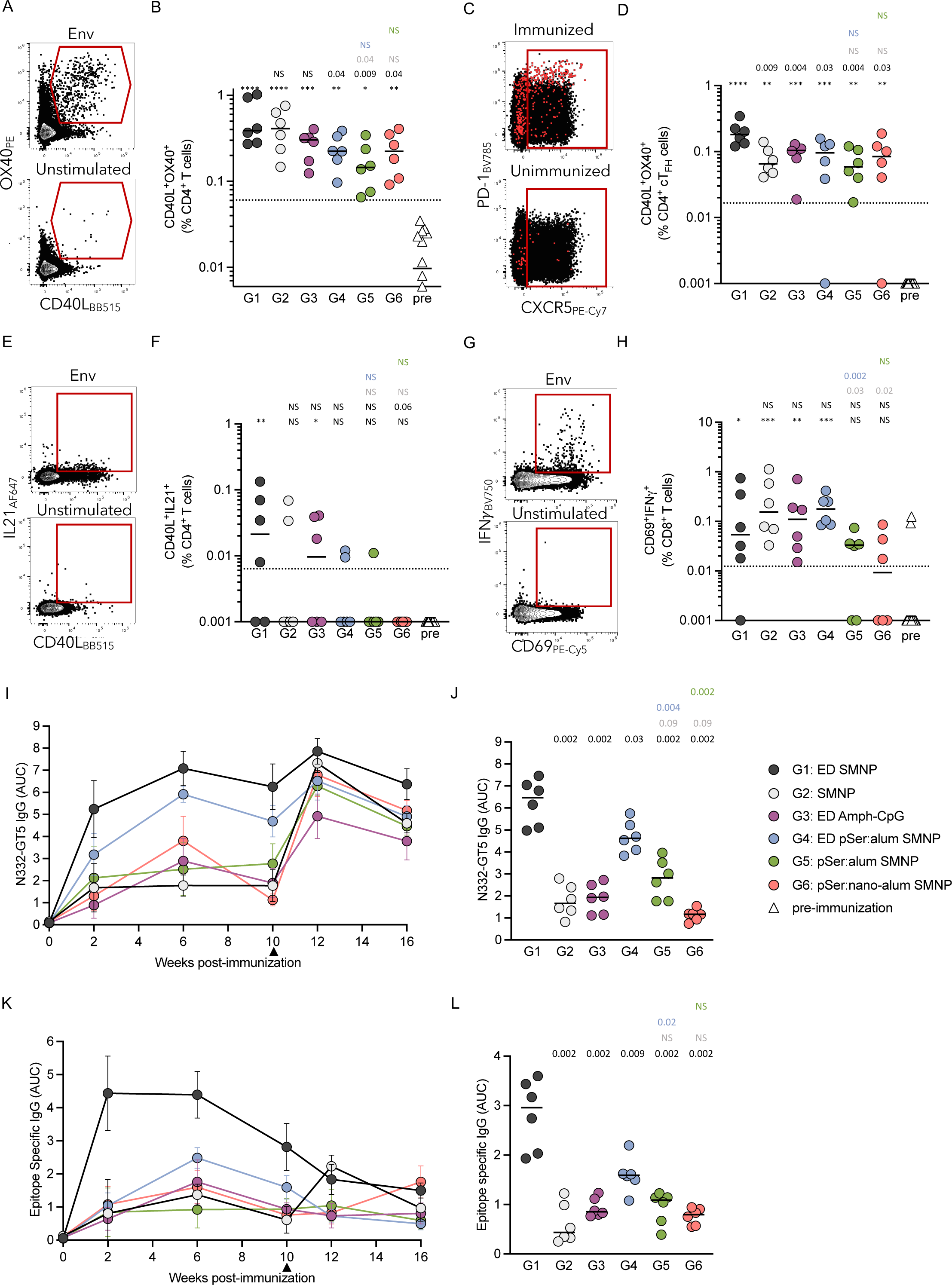
Vaccine-elicited T cell and serum IgG responses. **A)** Representative flow cytometry gating of an AIM assay performed at week 2 using N332-GT5 (Env) overlapping peptide pools or a DMSO control (unstimulated). **B)** Frequency of AIM^+^ (CD40L^+^OX40^+^) T cells out of total CD4^+^ T cells. **C)** Representative flow cytometry gating of an AIM assay performed at week 2 using N332-GT5 (Env) overlapping peptide pools. Red dots are Env+ cells. **D)** Frequency of AIM^+^ (CD40L^+^OX40^+^) cT_FH_ cells out of total CD4^+^ cT_FH_ cells. **E)** Representative flow cytometry gating of an AIM assay performed at week 2 using N332-GT5 (Env) overlapping peptide pools or a DMSO control (unstimulated). **F)** Frequency of AIM^+^ICS^+^ (CD40L^+^IL21^+^) T cells out of total CD4^+^ T cells. **G)** Representative flow cytometry gating of an AIM assay performed at week 2 using N332-GT5 overlapping peptide pools. **H)** Frequency of AIM^+^ICS^+^ (CD69^+^IFNg^+^) T cells out of total CD8^+^ T cells. **I)** Longitudinal area under the curve (AUC) of total antigen-specific serum IgG measured by ELISA. **J)** AUC of antigen-specific serum IgG at week 10 plotted by animal. **K)** Longitudinal area under the curve (AUC) of total epitope-specific serum IgG measured by ELISA. **L)** AUC of epitope-specific serum IgG at week 10 plotted by animal. Mean and SD are plotted in I and K, lines in all other graphs are median values. Pre-immunization time point included in all AIM/ICS assays. Statistical significance was tested using unpaired two-tailed Mann-Whitney tests with the p-values listed on each graph representative of the tests carried out, NS listed when p-value was >0.1. The row of asterixis in B, D, F, and H above the Y-axis indicate statistical significance for each group compared to the pre-immunizations samples using the following scale: NS > 0.05, * <0.05, ** <0.01, *** <0.001, **** <0.0001.

IL-21 ICS was performed to assess whether the vaccine-specific cells were able to produce IL-21, which is an important cytokine for B_GC_ and plasma cells (B_PC_) (AIM+ICS, CD40L^+^IL21^+^, **Fig. 4E**). Detectable IL21^+^ antigen-specific CD4 T cells were identified in some animals, with only G1 (4/6) and G3 (3/6) having significantly higher responses compared to pre-immunization (p = 0.001 (G1) and 0.013 (G3), **Fig. 4F**).

CD8^+^ T cell responses were examined by AIM+ICS to determine whether these protein immunizations primed antigen-specific CD8^+^ T cells (AIM+ICS, CD69^+^IFNγ^+^). All groups had at least 3 animals with detectable IFNγ-secreting CD8^+^ T cells, with all but G5 and G6 significantly higher than background (**Fig. 4G-H**).

Altogether, these T cell data show that all immunization strategies primed antigen-specific CD4^+^ T cells. cT_FH_ cell and IL-21 responses were particularly robust after ED SMNP immunization, consistent with the larger early GC responses seen in G1, which may have contributed to the strongest early recruitment of very rare BG18 Type 1 precursors in G1.

### Escalating dose with SMNP led to rapid induction of antigen-specific serum IgG

To further dissect the response to each delivery strategy and adjuvant combination, serum IgG titers were measured over time by ELISA, and AUC values were calculated (**Fig. 4I-L** and **Supp.Fig. 7A-B**). G1 had rapid induction of high titers of N332-GT5 specific serum IgG at week 2, which peaked at week 6 and remained high at the time of boosting (**Fig. 4I**). In contrast, G2 had low levels of serum IgG that remained stagnant after seroconversion at week 2 (**Fig. 4I**). Antigen-specific serum IgG peaked at week 6 in G3, G4, and G6, with declines at week 10. G1 maintained significantly higher titers at week 10 than all other groups (**Fig. 4I-J**). IgG AUC values for G1 were 5 times higher than G2 at week 10, demonstrating a significant enhancement in the ability of ED immunizations to induce high titers of antigen-specific IgG after priming compared to bolus immunization. Epitope-specific IgG AUC values in G1 were 6 times higher at week 10 compared to G2, further emphasizing the stark difference in the serum IgG response between ED and bolus priming (**Fig. 4K-L**. Median log10 endpoint titers G1 vs G2, 1.81 vs 0.39; **Supp.Fig. 7B**). G6 (nano-alum) exhibited the lowest IgG titers post-prime (**Fig. 4K-L**), contrasting with the relatively strong GC responses measured (**Fig. 2**). When post-prime antigen- and epitope-specific B_GC_ responses were graphed together with IgG AUC values at week 10, G6 is a clear outlier (**Supp.Fig. 7C-D**); exclusion of G6 increases the Pearson correlation values and significance of the B_GC_ cell to serum IgG association for both antigen- and epitope-specific responses (**Supp.Fig. 7C-D**). This suggests that although for most groups there is a strong association between GC and serum antibody responses, nano-alum immunization generates unusually divergent B_GC_ and B_PC_ responses. In total, the serum IgG data showed that early priming responses were much stronger after ED SMNP immunization compared to all other groups, with sustained high titers of antigen- and epitope-specific IgG in G1.

### Escalating dose primed larger B_mem_ responses but bolus immunization led to better memory recall responses post-boost

To evaluate the immune memory elicited by the different adjuvants and vaccine delivery mechanisms, antigen- and epitope-specific IgD^−^ memory B (B_mem_) cells were interrogated longitudinally by flow cytometry (**Fig. 5A, C**). G1 had the highest B_mem_ frequencies for both total antigen-specific and epitope-specific cells at week 6 (**Fig. 5B, D**). Most groups had modest measurable antigen- and epitope-specific B_mem_ cells at week 6 or 10 post-prime (**Fig. 5B, D, E, G**). At week 10, G1 had a significantly higher frequency of epitope-specific B_mem_ cells compared to all other groups and was 10-fold higher than G2 (**Fig. 5G**). Despite antigen-specific GC responses similar to G1 (**Fig. 2I**), G4 and G6 had significantly lower levels of antigen-specific B_mem_ at week 10 compared to G1 (**Fig. 5E, G**).

**Figure 5:**
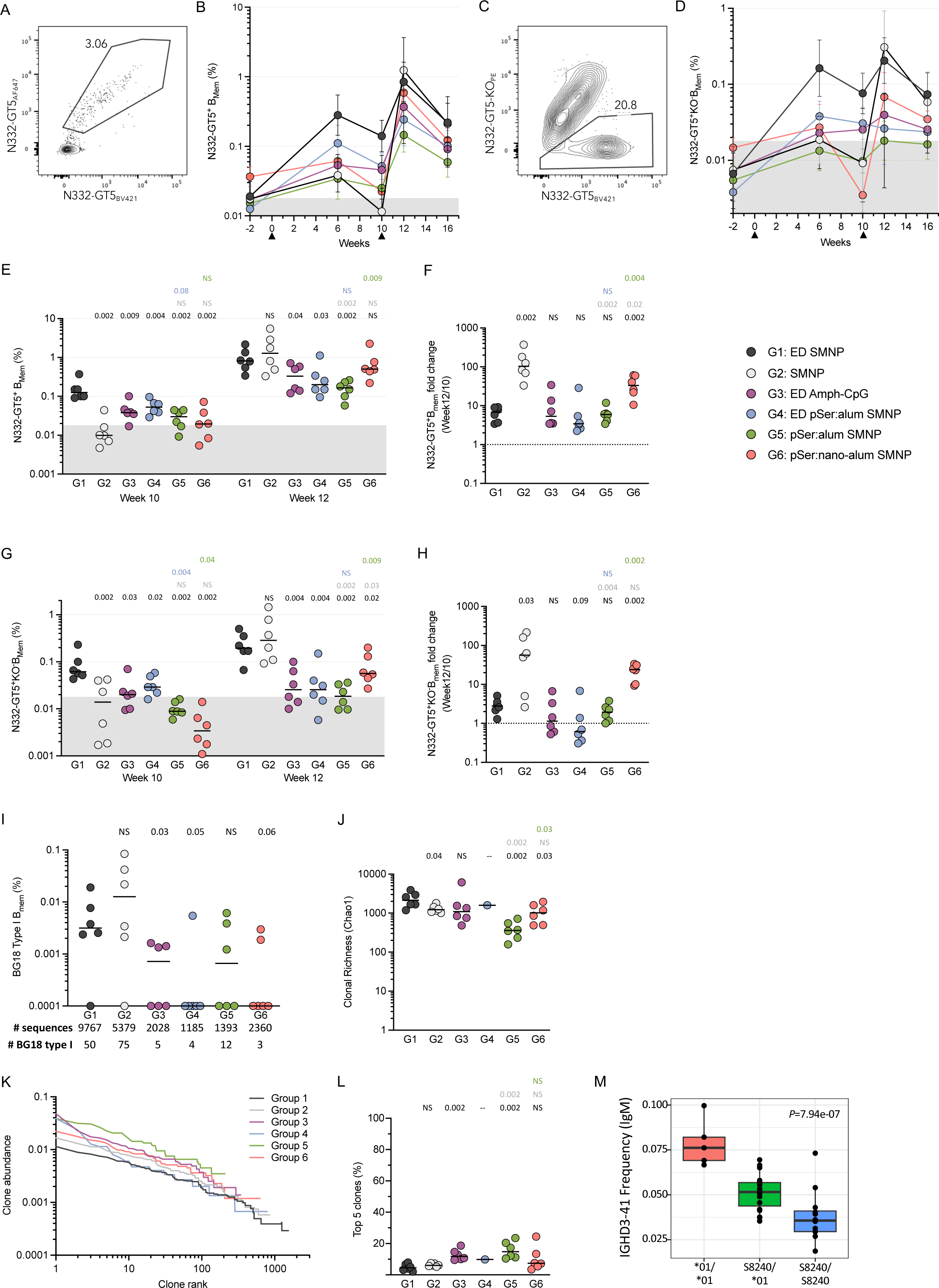
Antigen-specific. **B**_mem_ **responses. A)** Representative flow cytometry gating showing N332-GT5 antigen specific (N332-GT5-AF647^+^N332-GT5-BV421^+^) B_mem_ cells. **B)** Frequency of antigen-specific B_mem_ cells of total B cells from PBMCs over time for each group. **C)** Representative flow cytometry gating showing N332-GT5 epitope specific (N332-GT5-AF647^+^N332-GT5-BV421^+^N332-GT5KO-PE^−^) B_mem_. **D)** Frequency of epitope specific B_mem_ cells of total B cells over time for each group. **E)** Week 10 and week 12 antigen specific B_mem_ frequency plotted per animal. **F)** Fold change of week 12 antigen specific B_mem_ frequency over the week 10 antigen specific B_mem_ frequency. **G)** Week 10 and week 12 epitope specific B_mem_ frequency plotted per animal. **H)** Fold change of week 12 epitope specific B_mem_ frequency over the week 10 epitope specific B_mem_ frequency. **I)** Frequency of BG18 type I BCRs that are B_mem_ cells among total B cells plotted per animal at week 12. Numbers below indicate total number of paired BCR sequences recovered from each group and the total number of BG18 type I BCRs recovered. **J)** Clonal richness of the BCR sequences recovered from epitope specific B_mem_ cells from week 12 plotted as the Chao1 index on a per animal basis. Chao was calculated for all animals that had at least 1 sequence recovered. **K)** Clonal abundance curves for each group from sequences at week 12. Clonal abundance was calculated per animal then the mean abundance for each clone rank was plotted for each group. Animals with fewer than 5 total clones were excluded from abundance calculations. **L)** Cumulative abundance of the top 5 most abundant clones plotted per animal. The lines in all per animal graphs represent medians. **M)** Frequency of IGHD3-41 usage in IgM^+^ naïve B cells for 35 of the 36 animals plotted by specific genotype. Statistical significance was tested using unpaired two-tailed Mann-Whitney tests with the p-values listed on each graph representative of the tests carried out, NS listed when p-value was >0.1. Gray regions (B, D, E, G) represent B_mem_ LOD determined using pre-immunization samples.

After boosting at week 10, an increase in antigen-specific B_mem_ frequencies were observed for all groups. Unexpectedly, G2 exhibited a median antigen-specific B_mem_ frequency equal to G1 at week 12, despite having the lowest pre-boost antigen-specific B_mem_ frequency (**Fig. 5E**). The median epitope-specific B_mem_ frequency at week 12 was also equal between G2 and G1 (**Fig. 5G**). To explore this change further, the week 10 to 12 B_mem_ fold change was calculated for each animal (**Fig. 5F, H**). G2 had a 100-fold increase in B_mem_ (**Fig. 5F**), ~20 times greater than the increase observed for G1. Nano-alum G6 had the 2^nd^ lowest B_mem_ frequency post-prime, and exhibited a 30-fold increase at week 12 (**Fig. 5F**). Similar results were observed with epitope-specific B_mem_ (**Fig. 5H**). Despite major shifts in B_mem_ frequencies, the LN FNA antigen- and epitope-specific B_GC_ frequencies were mostly unchanged from week 10 to 12 (**Fig. 2E, H**). Notably, the G2 B_GC_ frequencies at week 12 were still 5- and 20-fold lower than G1 for antigen- and epitope-specific B_GC_ cells respectively, despite similar B_mem_ frequencies (**Fig. 2E, H**). In sum, ED immunizations formulated with SMNP led to the highest frequencies of antigen- and epitope-specific B_mem_ cells post-prime; however, booster immunization drastically increased the frequency of B_mem_ in response to bolus SMNP or nano-alum combined with SMNP, suggesting a substantial shift in the parameters regulating immunogenicity between the prime and the booster immunization.

### Bolus SMNP induced a high frequency of BG18 type I B_mem_ cells post-boost

Epitope-specific B_mem_ were sorted and sequenced from PBMCs at week 12. BG18 type I B_mem_ sequences were found in 5 of 6 animals in G1 (**Fig. 5I, Supp.Table 2**), consistent with the GC data (**Fig. 3A**) and results from the first study^25^. Unexpectedly, while no BG18 type I B_GC_ were detected in G2 at week 3, G2 had a high frequency of BG18 type I B_mem_ at week 12 that was not significantly different from G1 (**Fig. 5I**). The remaining study groups inconsistently developed BG18 type I B_mem_, which were only detected in 2 of 24 animals (**Fig. 5I**). Of note, despite robust GC responses in G4 and G6, only one animal from G4 and two from G6 had detectable BG18 type I B_mem_ (**Fig 5I**). The results observed for BG18_20-21AA_ and potential type III cells were similar to BG18 type I responses across the groups, with the caveat that G2 appeared to have more sporadic responses (**Supp. Fig. 8B-C**).

The clonal diversity of B_mem_ in the immunized animals was examined by multiple metrics. B_mem_ were more clonally rich and diverse across all groups compared to B_GC_, likely due to the inherently less clonal nature of circulating B_mem_ cells compared to B cells participating in a GC reaction in a single LN (**Fig 5J, Supp.Fig. 8D** compared to **Fig 3B-E, Supp.Fig. 5D**). G1 still had the greatest B_mem_ clonal diversity across multiple measures (**Fig 5J-L, Supp.Fig. 8D**). However, similar to Bmem frequencies above, G2 was much more similar to G1, with only clonal richness showing a significant difference between G1 and G2 (**Fig 5J**). Simpson diversity index and cumulative abundance curves showed similar trends (**Supp.Fig 8D, Fig 5K**). Immunodominance in immune memory, as assessed by the dominance of the top 5 B_mem_ clones, was lowest in G1, G2, and G6 (**Fig 5K-L**). Overall, the totality of the post-boost B_mem_ data showed that bolus or ED immunizations adjuvanted with only SMNP could prime rare BG18 type I precursors after N332-GT5 immunization. All other combinations inconsistently led to priming of BG18 type I precursors. In addition, the data indicated that, despite relatively low GC responses in G2 animals, a homologous booster immunization could generate highly expanded BG18 type I B_mem_ cells and a more clonally rich and diverse response than initially expected.

### IGHD3-41 Immunogenetics

BG18-like type I BCRs utilize the RM D gene *IGHD3-41*^25^. In both humans and RMs, there is extensive genetic diversity within the immunoglobulin (IG) gene loci, including coding and non-coding single nucleotide variants, as well as large structural variants that can result in changes in IG gene copy number ^30,31,38,52–55^. To date, >800 IG alleles have been characterized in RMs, including multiple alleles at the IGHD gene, *IGHD3-41*^30,38^. The coding sequence of IGH genes is known to be important for antigen binding and development of specific responses to pathogens^56^ and in the case of BG18, the D gene coding sequence is an important part of the paratope^12,25^. To assess the genetic compatibility of animals in this study for BG18 type I responses, we carried out long-read sequencing of the IGH locus in all 42 animals to genotype *IGHD3-41* alleles. All animals carried *IGHD3-41* in their genome and were determined to be either homozygous or heterozygous for the alleles *IGHD3-41*01* and *IGHD3-41*01_S8240* (**Supp.Table 3)**. Both of these alleles are capable of making BG18 type I responses using reading frame 1 or 3 (**Supp.Table 3**).

IG immunogenetic studies have shown that allelic variants can associate with differential gene usage in the naïve B cell repertoire, which has the potential to skew the composition of B cell and antibody responses^7,57–59^. For 35 animals, D gene usage in the IgM repertoire was examined using bulk adaptive immune receptor repertoire sequencing (AIRR-seq). Animals that were *\*01/*01* homozygotes used *IGHD3-41* at a higher frequency in naïve IgM^+^ B cells than animals that were *\*01/*01_S8240* heterozygotes or *\*01_S8240/*01_S8240* homozygotes (~1.5- and 2-fold higher, respectively. p=7.94×10^−7^, **Fig. 5M**. The three genotypes were relatively well distributed throughout the groups, with heterozygotes being the dominant genotype represented (**Supp.Fig. 9A-B**). When testing for associations between genotype and post-boost BG18 type I BCR frequencies, *\*01/*01* homozygotes had higher frequencies, but this association did not reach significance when looking at all animals together (**Supp.Fig. 9B-C**). Despite the relatively large number of total animals used in this study, it was still underpowered to detect differences between genotypes due primarily to the scarcity of *\*01/*01* individuals. Nevertheless, all three genotypes were capable of making BG18 type I responses after N332-GT5 vaccination (**Supp.Fig. 9B-C**), and the three genotypes were randomly distributed between the groups (**Supp.Fig. 9A-B**). Thus, the *IGHD3-41* genotypes influence on B cell precursor frequencies did not explain the majority of the inter-group BG18 type I B cell response differences observed.

### Serum IgG levels inversely predicted booster immunization outcomes

Post-boost serum IgG titers were quantified (**Fig. 4I, K**). G2 antigen-specific IgG increased 5-fold from week 10 to 12, while only a modest change was observed for G1 (**Fig. 6B, D**). At week 12, antigen- and epitope-specific IgG AUC values for G2 were similar to G1 (**Fig. 4I, K**. **Fig. 6A, C)**. A clear inverse relationship was observed for all groups between pre- and post-boost IgG titers, whereby the higher the week 10 IgG titers the smaller the fold increase after boosting (r = −0.96. P < 0.0001. **Fig. 6E**). A similar relationship was seen with epitope-specific IgG (r = −0.71. P < 0.0001. **Supp.Fig. 10A**). A significant inverse relationship was also seen between the B_mem_ fold changes (boost:pre-boost) and pre-boost serum IgG titers (week 10) across study groups (r=−0.52, p=0.001. **Fig. 6F**). These data indicate that very high levels of circulating N332-GT5 specific IgG at week 10 in G1 were associated with a blunted B cell recall response to homologous booster immunization, as observed in the modest increases in B_mem_ (**Fig. 5H**), BG18 type I BCRs (**Fig. 5I**), and serum IgG (**Fig. GD**). In contrast, the low levels of pre-boost IgG in G2 allowed for strong responses to the booster immunization, involving a dramatic 100-fold increase in antigen-specific B_mem_, detectable BG18 type I B_mem_ in most animals, and large increases in serum IgG titers.

**Figure 6:**
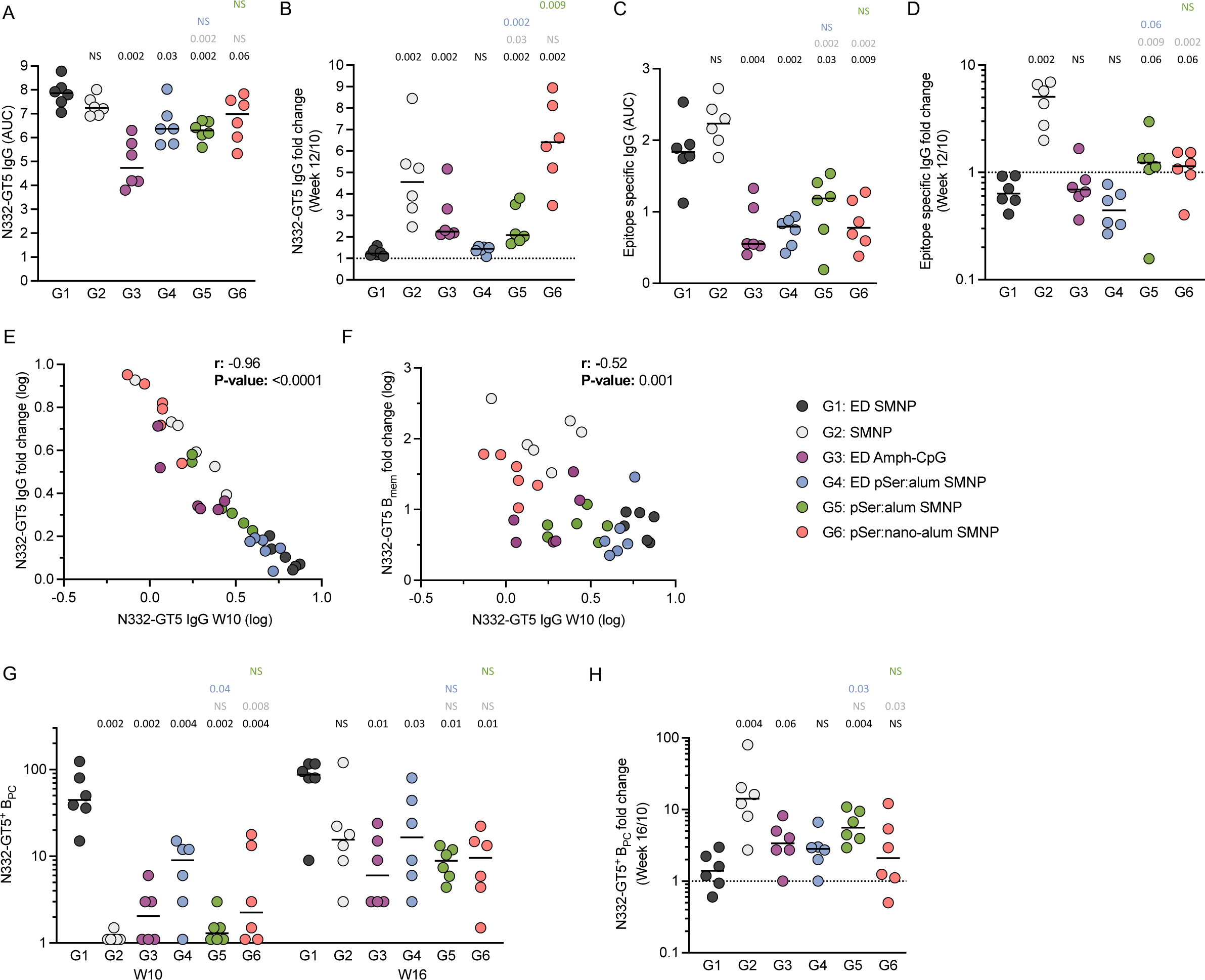
Level of antigen specific circulating IgG predict boost outcomes. **A)** AUC of antigen-specific serum IgG at week 12 plotted by animal. **B)** Fold change increase in serum IgG from week 10 to 12 calculated by taking week 12 AUC over week 10 AUC. **C)** AUC of epitope-specific serum IgG at week 12 plotted by animal. **D)** Fold change from week 10 to 12 for N332-GT5 epitope specific serum IgG. **E)** Correlation between fold change increase from week 10 to 12 and week 10 AUC. r and p-value are from pearson correlation analysis. **F)** Correlation between fold change increase for antigen-specific B_mem_ from week 10 to 12 and week 10 IgG AUC. r and p-value are from pearson correlation analysis. **G)** N332-GT5 specific BM-B_PC_ measured from bone marrow aspirates by ELISpot assay at weeks 10 and 16. **H)** Fold change from week 10 to 12 for BM-B_PC_. Statistical significance was tested using unpaired two-tailed Mann-Whitney tests with the p-values listed on each graph representative of the tests carried, NS listed when p-value was >0.1.

### Bone marrow plasma cells

The durability of the humoral immune response for each group was assessed by quantifying N332-GT5 specific bone marrow (BM) B_PC_ (**Fig. 6G, Supp.Fig. 10B-C**). Only G1 developed substantial antigen-specific BM B_PC_ after priming (week 10, **Fig. 6G**), with levels ~40-fold greater than G2. All groups showed an increase in BM B_PC_ 6-weeks after the booster immunization (week 16, **Fig. 6G-H**). G2 had the most dramatic increases from week 10 to 16 (**Fig. 6H**), which correlated with the strength of the IgG and B_mem_ response post-boost. G2 had the second highest BM B_PC_ responses post-boost, although still 5-fold lower than G1 (**Fig. 6G**). However, the ELISA data (**Fig. 4I, K**) suggests that in G2 much of the post-boost response is short-lived, while both antigen- and epitope-specific responses appear to be more stable in G1, evident by the more rapid decline in titers from week 12 to 16 for G2. The BM B_PC_ data reinforced that SMNP adjuvanted bolus or ED immunizations led to the strongest outcomes across all measures of humoral immune responses.

### BG18 type I BCRs had more SHM and higher affinity for boosting candidates

Similar levels of circulating BG18 type I B_mem_ cells were observed after booster immunization for both G1 and G2 (ED and bolus SMNP, **Fig. 5I**). BG18 type I B_GC_ cells were undetectable in G2 at week 3, and antigen-specific B_GC_ cells were 50-fold lower in G2 than G1 at week 3 (**Fig 3A and 2H**). Thus, the week 3 and week 12 BG18 type I cell outcomes were discordant. We therefore endeavored to better understand the underlying immunological processes resulting in these large changes. This topic was of particularly interest given the very rare precursor cells involved. Three plausible scenarios were considered. One possibility was that many fewer BG18 type I cells were primed in G2 compared to G1, but that post-boost those few BG18 type I clones in G2 massively expanded because of the high affinity interaction with N332-GT5 and the absence of competing circulating IgG. A second possible scenario was that BG18 type I cells were only primed in G2 after the second immunization, whereas BG18 type I cells were primed and expanded after the initial ED immunization of G1. A third possible scenario was that BG18 type I cells were primed in G2 after the first immunization, but at the 3-week time point G2 still had low vaccine-specific GC responses, which subsequently peaked 6-10 weeks post-prime (**Fig 2H, K**) resulting in the very rare BG18 type I cells being recruited into the GCs substantially later post-prime for G2 compared to G1, with a booster immunization required to expand the BG18 type I cells sufficiently to be detectable in G2. We resolved these three scenarios through a series of experiments, including assessment of clonality, quantitation of somatic hypermutation (SHM), and measurements of antibody affinities derived from G1 and G2 BG18 type I BCRs.

First, BCR sequence analysis revealed that responding B cells in both G1 and G2 had similar numbers of total BG18 type I clonal families (**Fig. 7A**). The clonal families had equally diverse usage of different IGHV genes (**Fig. 7B**). The clonal families also had equally diverse light chain usage in both groups (**Supp. Fig. 11A**). These data indicated scenario #1 was unlikely.

**Figure 7:**
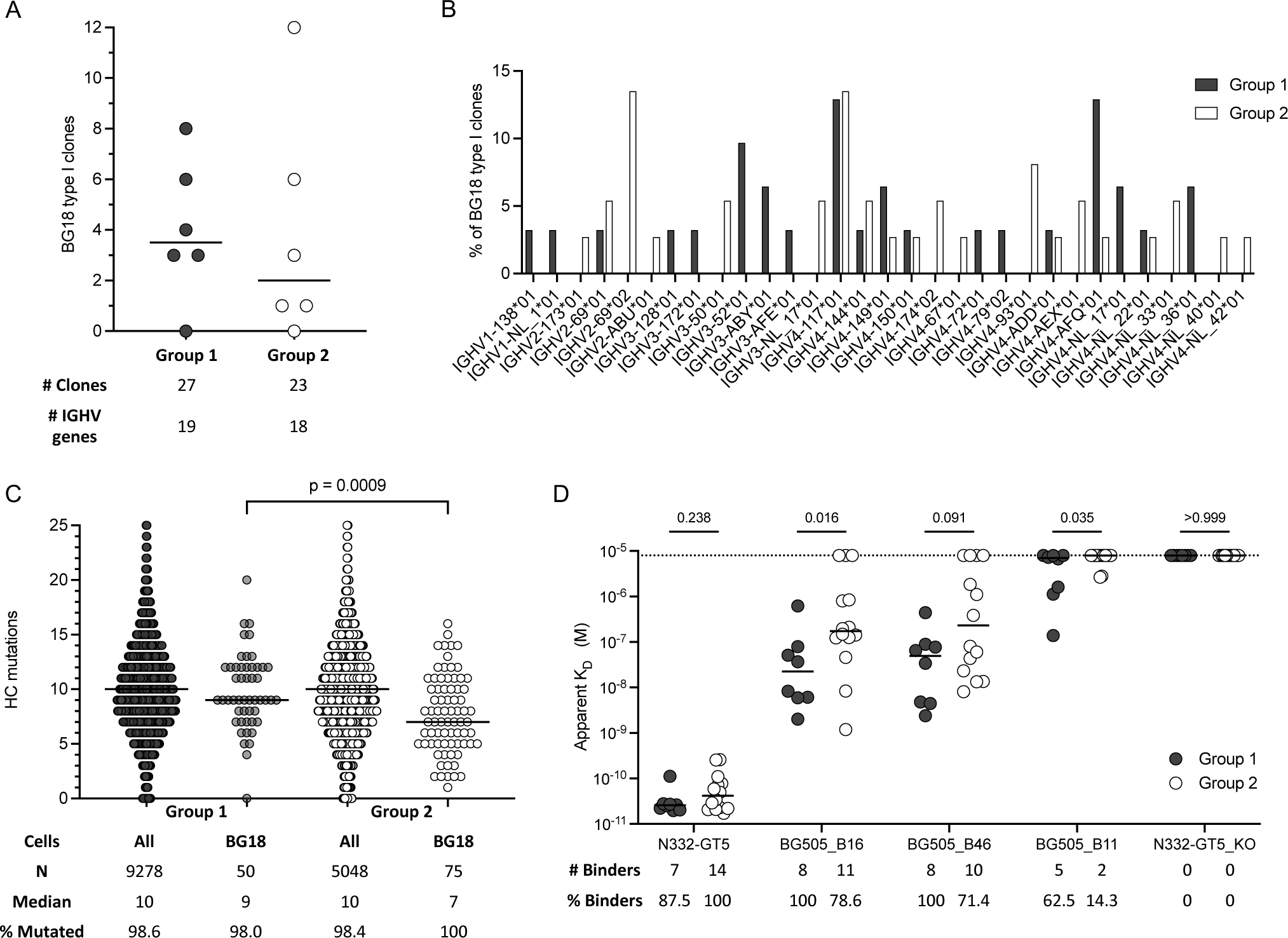
BG18 type I BCRs have more SHM and higher affinity for boosting candidates. **A)** Quantitation of BG18 clones in each group and the number of IGHV genes used by those clones. **B)** Graph of the percentage of BG18 type I clones for each group that use the specified IGHV genes. **C)** Heavy chain nucleotide mutations plotted on a per cell basis for each group. Both total recovered BCRs and BG18 type I BCRs are shown with the number of sequences in each, the median mutations listed, and the percent of all sequences with at least 1 mutation. Statistical significance was tested using unpaired two-tailed Mann-Whitney test. **D)** Binding affinities (Apparent K_D_) for a random subset of BG18 type I BCRs expressed from groups 1 and 2 to N332-GT5 and three potential boosting immunogens obtained by capturing IgG as the ligand and flowing trimers as analyte. Number and percent of binders are listed for each comparison (G1: n = 8, G2: n = 14). Dotted line represents the limit of detection for this assay. Statistical significance was tested between groups using unpaired two-tailed Mann-Whitney tests.

Scenario #2 (priming of BG18-like cells only after the G2 booster immunization) could be tested by examination of SHM between G1 and G2. In that scenario, BG18 type I BCRs from G2 B_mem_ at week 12 would have low or no SHM, whereas BG18 type I BCRs from G1 B_mem_ would have substantial SHM, while the total antigen-specific B_mem_ populations would have similar mutations between groups. Scenario #3 could also be resolved by SHM analysis, as cells experiencing less time in GCs would be expected to have less SHM. SHM was analyzed for all groups (**Supp. Fig. 11B-C**). Overall SHM heavy chain (HC) mutation counts of total antigen-specific B cells were similar between G1 and G2 (**Fig. 7C**), with > 98% of clones containing HC SHM in both groups, consistent with the vast majority of the B_mem_ derived from the antigen-specific B_GC_ responses observed in both groups post-prime (**Fig. 2H-I**). Notably, when BG18 type I BCRs were examined, > 98% of clones also contained HC SHM in both groups (**Fig. 7C**). G2 did have more BG18 type I BCRs that had 1-4 HC mutations at week 12 (**Fig. 7C**); however, all but one of these cells belonged to clonal families with BCRs that had >5 HC mutations, making it unlikely that they were primed just two weeks earlier (**Supp.Table 2**). These results suggested that scenario #2 was implausible to explain the majority of the BG18 response in G2. In contrast, when SHM of BG18 type I BCRs was quantitatively assessed, G2 had fewer HC mutations per BCR than G1 (p = 0.0009, **Fig. 7C**), while SHM of non-BG18 BCRs was indistinguishable between G1 and G2 (**Fig. 7C**). Thus, a parsimonious interpretation of the data is that BG18 type I B cells from G2 likely participated in GC reactions for a shorter period of time after the initial priming than did the BG18 type I cells from G1, based on SHM levels.

To examine the functional consequences of the SHM differences, a selection of BG18 type I BCRs from each group were produced as IgG antibodies, and binding affinities were determined. Antibodies from both G1 and G2 exhibited very high affinity for N332-GT5, and no significant difference was observed between the groups (**Fig. 7D**). All antibodies were epitope-specific, as expected, exhibiting no binding to N332-GT5KO (**Fig. 7D**). Affinity to three heterologous Env trimer boosting candidates^9,25^ was determined. 100% of BG18-like antibodies from G1 had detectable affinity for heterologous booster candidate BG505_B16, the most similar antigen to N332-GT5, while only 79% (11/14) of the antibodies from G2 bound BG505_B16. BG505_B16 affinities were significantly stronger for G1 antibodies compared to G2 (p = 0.016, **Fig. 7D**). Affinities were also significantly stronger for G1 antibodies compared to G2 for BG505_B11, the most native-like Env booster candidate tested (p = 0.035, **Fig. 7D**). Overall, these SHM and binding data consistently indicate that BG18 type I precursors from G2 were primed by the initial immunization but spent less time participating in the GC reaction compared to G1, with functional consequences observed by reduced binding affinity for heterologous boosting candidates.

## DISCUSSION

Despite their importance in generating quality immune responses to vaccination, the processes governing recruitment of rare bnAb-precursor B cells, and competition between rare and common B cell specificities, are only partially understood^6^. These processes have primarily been studied in mouse models using precisely controlled frequencies of adoptively transferred rare B cells^4,9–12,60,61^. Here we explored recruitment and differentiation of rare B cells in an outbred, non-transgenic, model using an HIV germline-targeting priming immunogen and demonstrate that different antigen delivery strategies and adjuvants can extensively impact the priming, expansion, differentiation, and affinity maturation of rare precursor B cells.

Despite all six groups inducing antigen-specific GC responses, only ED SMNP (G1) drove early recruitment of BG18 type I precursors detectable at week 3. After a booster immunization, BG18 type I B_mem_ cells were identified in at least one animal from every group, indicating that recruitment of these cells to GC reactions and subsequent differentiation to B_mem_ is possible under a variety of immunization conditions. Most strikingly, a priming bolus immunization with SMNP (G2) recruited BG18 type I B cells to GCs, despite very small GC reactions observable by week 3 post-prime, and these cells could be rapidly expanded upon boosting.

Kinetics of naive B cell recruitment may play a large role in the results seen here, particularly the early GC data. Finding BG18 type I cells in at least one animal in every group at week 12 indicates that some recruitment of these rare precursors can occur under all six conditions tested, but is inconsistent at best in most groups. The frequencies seen amongst the week 12 B_mem_ cells is likely more representative of the true post-prime recruitment efficiency for each group. It is possible that in the week 3 GCs there are BG18 type I cells present at such low frequencies that we could not yet detect them. Nevertheless, in the case of the bolus immunization group the epitope-specific B_GC_ data suggest that these cells may enter GCs later under those conditions. The SHM data also suggest that the BG18 type I cells from the bolus group have spent less time in the GC than those after ED immunization, which is consistent with later recruitment. Those outcomes are consistent with early T cell help likely playing a crucial role in the kinetics of early GC development and diverse B cell recruitment to GCs^38,62,63^. This was evidenced by the most robust early GC-T_FH_ and cT_FH_ responses being observed in G1, which may be driving many of the other outcomes.

We previously estimated that it was conceivable that ~1.2 × 10^9^ naïve B cells could be “screened” in LNs for antigen-recognition under relatively optimized immunization conditions in humans^6^. We can extend this calculation, using similar assumptions, to RMs using updated data. There are ~5 million B cells per RM inguinal LN. We, and others, have shown that multiple LNs in a drainage region will uptake antigen^39,64–67^ and that after ED immunization with SMNP this may be 3-5 LNs in a specified region^39^ (~20 million B cells). It has also now been shown that after SMNP immunization the antigen drains to deeper LN clusters, probably tripling the number of LNs engaged (~60 million B cells)^68^. The immunizations in this study were done bilaterally, doubling the number of LNs engaged (~120 million B cells). Naïve B cells are constantly moving through lymphatics, and in humans exhibit a LN dwell time of 10-16 hours^69–71^. After a bolus immunization it is reasonable to assume that naïve B cells can be recruited during the first 48-hours^72^, which would perhaps represent 4x turnover of the naive B cell population during that timeframe. The data suggest that after ED immunization naïve B cells may be recruited for 2-to 3-weeks^37,38,73^, which plausibly represents 28 to 42x turnover of the naive B cell population per LN during that 2-to 3-weeks. Furthermore, influx of cells into LNs after immunization can lead to swelling and 2-to 4-fold increased cell numbers^72^. Those estimations result in a calculated number of ~13 × 10^9^ naive B cells potentially being “screened” *in vivo* for antigen binding after ED SMNP immunization in RMs.

Although we do not know the true number of BG18 type I precursors that can bind to and be activated by N332-GT5 in RMs, in humans, BG18 type I precursors were found at ~1 in 50 million naïve B cells^12,25^. Based on previous precursor frequency calculations that number is likely to be 2-8-fold lower in RMs (1 in 100-400 million^25^). Taken together, these calculations would indicate that between 32 and 130 BG18 type I naïve B cells are potentially screened for antigen binding in LNs after N332-GT5 ED immunization with SMNP in RMs.

Experimentally, we observed that in G1 that there was a median of 3.5 BG18 type I clonal families per animal (range 0-8, **Fig. 7A**) in the B_mem_ compartment. Using rarefaction analysis on the week 12 B_mem_ sequencing results, we estimated that ~50% (range of 40-75%) of the B_mem_ repertoire was sampled in the animals from G1 (**Supp. Fig. 7E**). This indicated that the true number of BG18 type I clonal families present in each animal was likely double that observed experimentally (median 7, range 0-16). These calculations were undertaken using assumptions based on experience with ED immunization in RMs, but the numbers are likely much lower in non-SMNP and bolus immunization animals, which underscores the difficulty in eliciting these types of rare responses in NHPs and likely humans as well.

When pSer modification was used to deliver a native HIV Env trimer in mice, increases in both antigen-specific GC B cells and IgG titers were seen compared to normal bolus immunization^43^. The same pSer trimer adjuvanted with SMNP led to slightly better total humoral responses, including development of autologous nAb titers, in NHPs, associated with the its extended release and valency attributes of pSer ^45^. It was therefore unexpected that bolus immunization led to better priming of BG18 type I cells compared to pSer:alum SMNP herein. Even more unexpected was the poor recruitment of BG18 type I cells seen when delivering pSer modified N332-GT5 by ED. This group had some of the highest GC responses, but those large numbers of B_GC_ did not translate into efficient recruitment of rare BG18-like precursors. Despite this, the nano-alum group exhibited unique immunological characteristics that suggest that nano-alum may have a distinct mechanism of action compared to traditional alum and further study is warranted.

The other adjuvant tested, Amph-CpG, performed worse than SMNP in ED immunization. Amph-CpG has shown promise in modulating immune responses against SARS-CoV-2 vaccines in NHPs^50^. Further work is needed to elucidate how engaging different TLRs leads to differences in adjuvanticity compared to SMNP.

One of the more striking outcomes presented here was the large boosting response seen for the bolus immunization group, which inversely correlated with the circulating IgG titers pre-boost. The outcome implicated antibody feedback as a major factor in the booster immunization outcomes. Antibody feedback has recently seen a resurgence in research, in part due to observations surrounding antibody responses to COVID-19 vaccination^74–77^. Antibody feedback can exert influence through multiple mechanisms. One possibility is that the high level of circulating IgG leads to antigen clearance effectively lowering the dose of the boosting immunization. Antigen clearance traditionally occurs through Fc receptor mediated uptake of immune complexes by non-follicular antigen-presenting cells^77^. Another possibility was epitope masking by IgG produced by BG18 type I B cells. In a recent knock-in mouse study of N332-GT5 immunization, the authors found that excess antigen could overcome the antibody-dependent clearance of antigen by effectively depleting the circulating IgG and allow for better secondary GC responses^78^. Others have shown that IgG to dominant epitopes can lead to better responses to sub-dominant epitopes in subsequent immunizations by focusing the GC response^74,76,79–82^. ED or other slow delivery immunization strategies likely enhance priming of more diverse B cell responses due to engagement of this mechanism involving the earliest antibodies^37,38,73^. It is conceivable that in the bolus immunization group, the low circulating IgG titers of off-target but antigen-specific IgG could lead to immune complex formation and better uptake by follicular dendritic cells after booster immunization, while the lack of epitope-specific IgG allows for a focusing of the response on epitope^83^. Whereas after ED, the extremely high titers of antigen- and epitope-specific IgG led to both antigen clearance and masking of the N332 epitope on residual antigen. In the pSer:alum ED group, the epitope-specific response may have been low enough to minimize any epitope masking, but it is conceivable that the total IgG titers were sufficiently high as to lead to antigen clearance. It remains an interesting observation that although the circulating titers at week 10 predict the post-boost B_mem_ frequencies, there is no correlation with the antigen- and epitope-specific B_GC_ frequencies from week 10 to 12.

In mice, many vaccination strategies induce high titers of circulating antibodies after a single immunization, which allows antibody feedback to be a potentially major factor^77,78,80,82^. It would be expected that the impact of antibody feedback would be more modest in contexts of lower antibody responses. In large animal models and humans, single immunizations most often generate relatively low antibody responses, which are also often transient^38,84–86^. The experiments described herein include three features resulting in high antibody titers after the priming immunization. One feature is the nature of the antigen itself, which is highly immunogenic due in part to a substantial glycan hole^9,19,25,29^. Two additional features are the use of ED immunization and SMNP, which both enhance rapid induction of antibody responses. These features make this model system have characteristics wherein antibody feedback mechanisms may be more likely observed, compared to contexts where priming elicits low antibody titers.

We previously found that combining pSer:alum delivery and SMNP augmented induction of autologous neutralizing responses in RMs to native-like HIV trimers over either vaccine strategy alone^45^, in contrast to the present study where pSer:alum delivery did not improve the recruitment of rare BG18-like precursors. An important difference in these two scenarios, which may play an important role in the B cell response, is epitope availability. The early antibody response to soluble native-like Env trimers if dominantly focused on the irrelevant base of the immunogen ^37,38,42^. Vaccine strategies like ED that amplify the early antibody response against this epitope may promote capture of antigen on FDCs and allow more time for rare precursors targeting other sites to be recruited to a GC. By contrast, N332-GT5 Env trimer is engineered with a prominent glycan hole around the target BG18 epitope, relatively distant from the Env base, and adoptive transfer models of rare precursor B cell recruitment by this immunogen have shown that the Env trimer base is no longer dominant in this scenario, but rather B cells binding in the vicinity of the new glycan hole epitope instead dominate^12^. In this situation, antibody feedback may hinder rather than help the development of the desired on-target B cell response.

The comparison between ED and bolus immunization here is additionally valuable as it informs ongoing clinical development of N332-GT5 as an HIV vaccine candidate. Currently, the first-in-human clinical trial for both N332-GT5 and SMNP is ongoing (ClinicalTrials.gov Identifier: NCT06033209). In that study, ED and bolus immunizations with N332-GT5 + SMNP are being compared head-to-head. The results presented here will help inform the analysis of that trial and provide a benchmark for comparing the priming efficiency in humans. Indeed, our results suggest that bolus immunization with suitably designed germline-targeting trimer immunogens and appropriate adjuvants may suffice for inducing bnAb precursor responses in humans. These data demonstrate that adjuvant selection is key, and the capacity of SMNP to induce large GC responses recruiting highly diverse B cells, including very rare precursors, will be an important component of any protein-based germline-targeting immunization regimen.

The next step for development of a germline-targeting vaccine to induce bnAbs will rely on shepherding of the primed precursors towards bnAbs, and therefore the BG18-class B cells induced by N332-GT5 must be able to engage more native-like HIV Env trimers, without being blocked by serum antibodies. At this time, the *in vivo* affinity threshold for boosting the primed BG18 type I responses in NHPs is not clear, but knock-in mouse data suggests that even modest affinities can lead to robust boosting^9^. We hypothesize that BG18 type I cells after ED or bolus SMNP + N332-GT5 have sufficient affinity to the tested immunogens as to be boosted *in vivo* by select heterologous Env trimers. Notably, ED+SMNP priming elicited B cells with higher affinity for booster immunogen candidates, suggesting that approach may allow for earlier introduction of native-like antigens in the immunization sequence.

In sum, here we demonstrated that both ED and bolus immunization can lead to priming and expansion of rare BG18 type I B cells in NHPs when adjuvanted with SMNP and represent the first direct study of rare B cell recruitment under multiple conditions in an outbred, non-transgenic, animal model. In the context of germline-targeting immunizations, the quality of the B cells was higher in the ED group, consistent with more rapid recruitment of B cells to GCs and more extensive SHM in GCs, along with differential regulation of the rare B cell responses by the different concentrations of circulating IgG generated by the different immunization conditions.

## STAR METHODS

### Adjuvant production and formulation

Alhydrogel aluminum hydroxide adjuvant was purchased from Invivogen and used as received. SMNP adjuvant was synthesized and characterized as previously described^39^. Amph-CpG consisting of the class B CpG 7909 sequence (5’-TCG TCG TTT TGT CGT TTT GTC GTT-3’) conjugated at the 5’ end to a diacyl lipid tail (ref ^87^)was prepared by solid phase synthesis by Axo Labs.

Nano-alum was generated by dispersing aluminum hydroxide adjuvant in the presence of a phosphate-containing PEG-lipid surfactant. Alhydrogel (10 mg, Invivogen) was diluted to 1 mg/mL in deionized water and combined with 10 mg of 1,2-distearoyl-sn-glycero-3-phosphoethanolamine-N- [amino(polyethylene glycol (PEG))-2000] (Avanti Polar Lipids). The mixture was transferred to a 20 mL glass scintillation vial and cooled to 4°C in ice water. While still on ice, the mixture was sonicated with a Misonix Microson Ultrasonic Cell Disruptor XL2000 probe tip sonicator for 30 minutes, during which it transitioned from opaque to opalescent. Care was taken to position the sonication probe deep enough in the mixture to prevent foaming. Output power at the start of the sonication was set to ~20, with the power turned down as sonication progressed and the solution transitioned from opaque to translucent. To remove excess alum aggregates, the sonicated material was then incubated at 25°C for five days to enable precipitation of any residual non-stabilized alum. The supernatant containing the suspended nano-alum was collected, and nanoparticle formation was verified via dynamic light scattering on a Malvern Zetasizer Nano instrument. Nano-alum used in this study was measured to have a Z-average diameter of ~70 nm and a low PDI.

### Protein production and pSer modification

For the expression of the N332-GT5 immunogen, a stable CHO cell line clone was developed utilizing the Leap-In transposon expression system (ATUM, US10041077). The DNA sequence encoding the N332-GT5 trimer and its corresponding signal peptide was first synthesized. Subsequently, expression constructs were designed and engineered based on the proprietary Leap-In transposon platform. The production of the trimer protein involved comprehensive upstream cell culture and downstream purification process development, followed by scale-up to large-scale cGMP manufacturing (details to be published in a forthcoming article). With the exception of the signal peptide, the amino acid sequence of this trimer, matches the N332-GT5 protein that was reported previously^25^.

The N332-GT5 trimer used for pSer conjugation contained the C-terminal linker sequence “GTKKKC”, previously optimized for Env trimers and referred to as nohis8^43^, placing a free terminal cysteine for subsequent pSer peptide coupling. The trimer was co-transfected with furin into 293F cells and purified using Galanthus nivalis lectin affinity chromatography (Vector Laboratories) followed by SEC purification using Superdex 200 16/600 PG column (Cytiva).

Trimers that were used as SPR reagents were expressed with C-terminal 6xHis-tag and purified using a HIS-TRAP column followed by size exclusion chromatography (SEC) on a Superdex 200 Increase 10/300 column (Cytiva) as described previously^25^. N332-GT5 and N332-GT5-KO trimers that were used as biotinylated sorting reagents were expressed with a C-terminal His-tag and avi-tag, purified using a HIS-TRAP column followed by SEC and biotinylated using a BirA biotin-protein ligase reaction kit (Avidity, catalog no. BirA500) as described previously^25^. Compared to the immunogens, the N332-GT5 sorting probes lacked the 241 and 289 glycosylation sites.

N332-GT5 trimers with a C-terminal Cys were conjugated with maleimide-functionalized pSer peptide tags and characterized as previously described^43^. Briefly, peptides carrying 4 phosphoserines and N-terminally functionalized with a maleimide group (mal-pSer_4_) were produced by solid phase synthesis, purified by HPLC, and their mass confirmed by matrix-assisted laser desorption/ionization-time of flight mass spectrometry. N332-GT5-Cys trimers were gently reduced with tris(2-carbox-yethyl)phosphine then reacted with 5 molar equivalents of mal-pSer4 in tris-buffered saline, followed by centrifugal filtration to remove unconjugated peptide. The degree of modification of trimers was quantified using a Malachite Green Phosphoprotein Phosphate Estimation Assay Kit (Thermo Scientific). Alum binding of pSer-conjugated N332-GT5 was assessed by labeling the trimer with an AlexaFluor dye and assessing bound protein following mixing with alum (loading) or following incubation of loaded alum with 10% mouse serum in tris-buffered saline for 24 hr as previously described^43^. Antigenicity profiles of alum-bound pSer-N332-GT5 were assessed by capturing alum particles on ELISA plates followed by addition of pSer-trimer to allow binding, washing, and then probing with indicated dilutions of monoclonal antibodies as previously described^42^.

### Mouse immunogenicity testing of nano-alum

All mouse studies were carried out at MIT under an IACUC-approved protocol following federal, state, and local guidelines. Female BALB/c mice (Jackson Laboratory) 6-8 weeks of age were immunized s.c. bilaterally at the tail base with alum/pSer-trimer or nano-alum/pSer-trimer. For flow cytometry analysis, lymph nodes were harvested, crushed to form a single-cell suspension, and subjected to staining for viability (Zombie Aqua Fixable Viability Kit, BioLegend), CD4 (BV711, BioLegend, RM4-5 clone), B220 (PE-Cy7, BioLegend RA3-6B2 clone), CD38 (FITC, BioLegend 90 clone), CXCR5 (PE, BioLegend L138D7 clone), PD-1 (BV421, BioLegend 29F.1A12 clone), and GL7 (PerCP-Cy5.5, BioLegend GL7 clone). Antigen-specific staining was done using biotinylated trimers conjugated to either streptavidin-BV605 (BioLegend) or streptavidin-APC (BioLegend). Analysis was performed on a Becton-Dickinson Symphony flow cytometer, and the resulting data were processed using FlowJo software. Anti-trimer serum Ig titers were assessed by ELISA. 96-well plates (Costar Corning) were coated overnight at 4°C with 1 μg/ml (50 μl/well) of streptavidin in PBS. Coated plates were blocked with 2% BSA in PBS, and incubated overnight at 4°C. Plates were then washed and incubated with 1 μg/ml biotinylated trimer in blocking buffer (2% BSA in PBS) for 2 h at 25°C, following which serial dilutions of serum samples were added to the plates. After 2 h incubation, plates were washed and incubated with HRP-conjugated anti-mouse IgG diluted 1:5,000 in blocking buffer. The reaction mixture was incubated at 25°C for 30 min. Plates were washed and treated with 50 μL TMB substrate for 10 minutes followed by addition 1M H_2_SO_4_ stop solution. Optical densities were read on a microplate reader at 450 nm with background correction at 540 nm. Endpoint serum titers were determined as the reciprocal of the highest serum dilution leading to a signal 0.1 OD units above background.

### Non-human Primates and Study Design

Adult Indian rhesus macaques (Macaca mulatta) were housed at the Emory National Primate Research Center and maintained in accordance with NIH guidelines. All procedures were approved by the Emory University Institutional Animal Care and Use Committee (IACUC) under protocols 202100128 and 201700666. Animal care facilities are accredited by the U.S. Department of Agriculture (USDA) and the Association for Assessment and Accreditation of Laboratory Animal Care (AALAC) International. RMs for this study were of mixed sex, an average age of 5.5 years, and a weight range of 4-7.2kgs at the time of the priming immunization. Animals were pair housed for the duration of the study. Thirty-six NHPs were assigned to one of six experimental groups—each group was comprised of 6 RMs. The immunizations were as follows: Group 1 (ED SMNP), Group 2 (SMNP), Group 3 (ED Amph-CpG), Group 4 (ED pSer:alum SMNP), Group 5 (pSer:alum SMNP), and Group 6 (pSer:nano-alum SMNP). Immunizations were administered subcutaneously (s.c.) in the left and right mid-thighs. SMNP was dosed at 375mg per injection site for all SMNP-containing groups. All alum-containing adjuvant (alum and Nanoalum) were dosed at 500mg per injection site. This is the alum dose that has been previously tested in pSer-based HIV Env immunizations in mice and RMs^43,45^. The Amph-CpG was administered at a dose of 2.5mg per injection site which was the optimal dose determined from a previous RM immunization experiment using SARS-CoV-2 RBD base protein immunogen^50^.

Veterinarians performed fine need aspirates (FNAs) to bilaterally sample lymph nodes (LNs) in the RMs. Palpation was used to identify draining LNs. A 22-gauge needle with an attached 3 ml syringe was passed into the LN a maximum of five times. Samples were then placed into RPMI media with 10% FBS and 1% Penicillin/Streptomycin. If red blood cell contamination was observed, an Ammonium Chloride-Potassium lysing buffer was used. Samples were counted, frozen down, and stored in liquid nitrogen until the time of analysis. Blood was collected throughout the study in NaCitrate CPT tubes (BD biosciences) for peripheral blood mononuclear cells (PBMCs) and plasma isolation and frozen. Serum was collected via serum clot tubes and frozen. Bone marrow aspirates were collected in heparin coated tube and analyzed via ELISPOT fresh. Animals were treated with anesthesia (ketamine 5-10 mg/kg or telazol 3-6 mg/kg) and analgesics for s.c. immunizations, LN FNA, bone marrow aspirates, and blood draws as per veterinarian recommendations and IACUC approved protocols. After completion of the proposed study and approval of the vet staff, animals were released to the center for reuse by other researchers.

### ELISA analysis of serum IgG titers

Serum IgG titers were measured by ELISA. Nunc MaxiSorp plates were coated with streptavidin 18 hr at 4°C and blocked with 2% BSA in PBS. Blocked plates were washed with washing buffer (PBS with 0.05% Tween-20) and incubated with Avi-tagged biotinylated antigen 18 hr at 4°C. Plates were washed again and serial dilutions of sera were added to plates and incubated for 2 hr at 25°C, following which the plates were washed and incubated with anti-rhesus IgG-HRP (BioRad). ELISA plates were developed with TMB substrate and stopped by sulfuric acid 2 M. Absorbance values at 450 nm were measured on a Tecan Infinite 200PRO plate reader, with a background correction wavelength of 540 nm.

### Immunoglobulin DNA and repertoire sequencing (AIRR-Seq) sample preparation

For immunoglobulin sequencing (AIRR-Seq), PBMCs were isolated from 15-20 ml of blood from each animal. For immunoglobulin repertoire sequencing. Approximately 5 M PBMCs were lysed in QIAGEN RLT buffer + 1% BME and stored at −80 C until RNA was purified using QIAGEN RNeasy kits. For sequencing of the immunoglobulin DNA loci, 5 M PBMCs were snap frozen and stored at −80 C.

### IGHD3-41 Genotyping

To genotype IGHD3-3 (“IGHD3.41”), we utilized a custom genomic IG targeted long-read Pacific Biosciences single molecule real-time (SMRT) sequencing method (KAPA HyperExplore, Roche), following previously published protocols^25,54,61,88^ [1-4]. Briefly, genomic DNA was isolated from peripheral blood mononuclear cells (PBMCs) collected from each animal using the DNeasy kit (Qiagen). DNA (~2.5 ug) was sheared with g-tubes (Covaris); size selected >5-6 Kb (Blue Pippin/Pippin HT, Sage Science); End Repaired and A-tailed (KAPA sequencing library protocol, Roche); and finally, sample-specific sequence barcodes and universal primers were ligated. Initial PCR amplification (8-9 cycles) was conducted using PrimeSTAR GXL polymerase (Takara), followed by size-selection and purification of amplified fragments 0.7X AMPure PB beads (Pacific Biosciences). Target-enrichment was performed by hybridization of custom IGH/K/L locus-specific oligonucleotide probes (KAPA HyperExplore, Roche). Oligo-bound fragments were recovered using streptavidin beads (Roche), and additional PCR amplification (16-18 cycles) were performed using PrimeSTAR GXL (Takara). Long-read sequencing libraries were prepared using the SMRTbell Express Template Preparation Kit 2.0 (Pacific Biosciences). Resulting libraries were sequenced on the Sequel IIe system.

HiFi reads for each animal were mapped to the RheMac10 reference genome (Mmul_10) using minimap2^89^. Phased single nucleotide variants representing two distinct alleles were resolved from HiFi reads spanning the IGHD3-3 gene (chr7: 168251081-168251167). At least 10 representative HiFi reads were required to include a given allele in the genotype of an animal.

### AIRR-Seq library preparation

The library preparation method was based on the protocol provided by Dr. Daniel Douek, NIAID/VRC^90,91^ and the IgM constant region primer was based on Corcoran et al, 2016^92^ and were described previously in Steichen et al, 2024^25^. RNA was extracted from sorted naive B cells in 350μL QIAGEN RLT buffer using the RNeasy Micro-DNase Digest protocol (QIAGEN) on QIAcube automation platforms (Valencia, CA). Clontech SMARTer cDNA template switching was used to perform the reverse transcription step. 8μL of RNA was mixed with 1 μL of 12μM 5’ CDS oligo(dT) primer for 3 seconds and incubated for 3 minutes at 72°C followed by at least 1 minute at 4°C. 8.5 μL of master mix comprising of 3.5 μL of 5x RT Buffer (250 mM Tris-HCl (pH 8.3), 375 mM KCl, 30 mM MgCl2), 1 μL Dithiothreitol, DTT (20 mM),1 μL dNTP Mix (10 mM), 1 μL RNAse Out (40U/µL), 1 μL SMARTer II A Oligo (12 µM), 1 μL Superscript II RT (200U/µL) was added to each reaction. After sealing and spinning the plates briefly, they were incubated for 90 minutes at 42°C and 10 minutes at 70°C. AMPure XP beads were used for purification of the first strand cDNA. For amplification of the rearranged BCR sequences, 19.3 μL of cDNA was mixed with 30.7 μL of master mix (25 μL of 2x KAPA Real-Time Library Amplification Kit (catalog# KK2702), 0.7 μL of 5PIIA (12 µM): forward primer and 5 μL of IgM primer (2 µM): reverse primer). The plates were sealed, vortexed for 5 seconds and centrifuged at 2000 RCF for 1 minute. Real-time PCR machine was used to visualize the amplification and the reaction was terminated in the exponential phase. The amplified BCRs were purified using the AMPure XP beads. Two more rounds of PCR amplification were performed for addition of barcodes and adapters. In the first round, 46 μL of master mix (25 μL of KAPA HotStart ReadyMix (catalog# KK2602), 2.5 μL of SYBR Green 1:10K and 18.5 μL of Nuclease-free water), 1 μL of P5_Seq BC_XX 5PIIA (10 µM), 1 μL of P7_i7_XX IgM (10 µM) and 2 μL of 1:10 diluted amplified rearranged BCR from previous PCR were mixed, sealed, vortexed and centrifuged. Real-time PCR amplification was performed and the resulting library was purified with AMPure XP beads. The last round of PCR amplification was performed for addition of the P5_Graft P5_Seq. Real-time PCR was carried out by mixing 44 μL of master mix (25 μL of KAPA HotStart ReadyMix (catalog# KK2602), 1 μL of P5_Graft P5_seq (10 µM): forward primer and 18 μL of Nuclease-free water), 1 μL of P7_i7_XX IgM (10 µM) (reverse primer) and 5 μL of amplified library from previous PCR followed by a final round of purification with AMPure XP beads. Agilent Bioanalyzer was used to assess the quality of the libraries and after pooling the libraries were sequenced on an Illumina MiSeq as 309 base paired-end runs. Primers used for AIRR-Seq library preparation and sequencing are listed below.

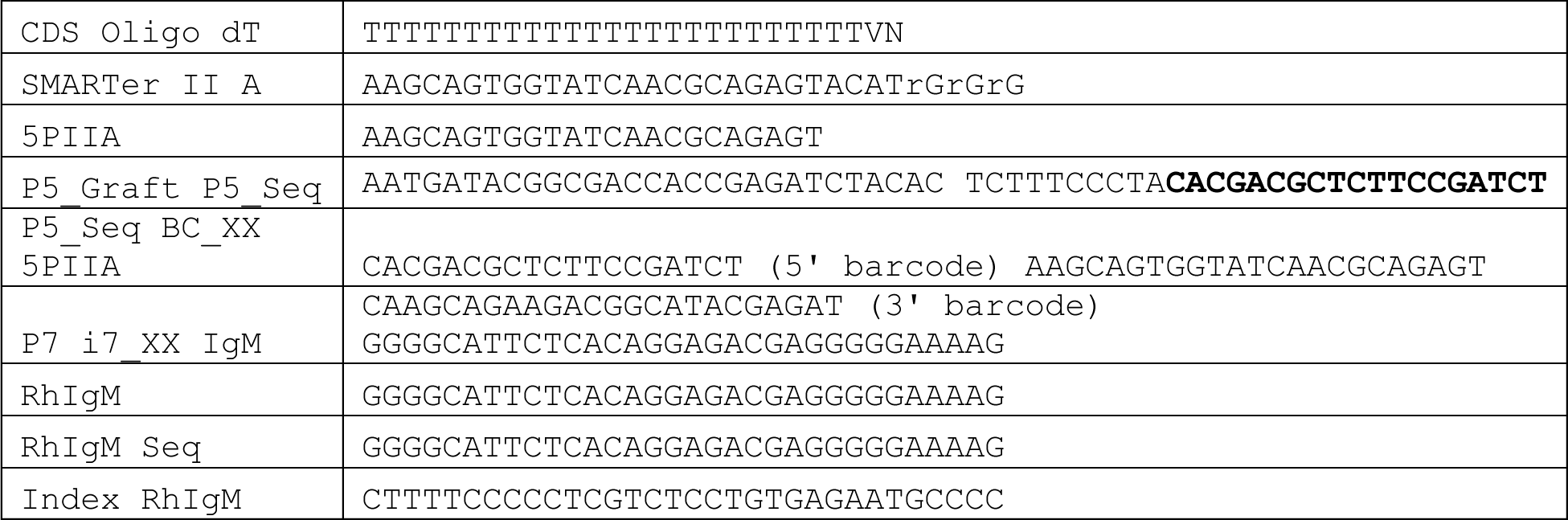

### AIRR-Seq data analysis

Fastq files were demultiplexed using a custom script that searches for an exact match and trims the 5’ barcode followed by the 5’ RACE primer sequence with the corresponding number of upstream random nucleotides. This script uses seqtk v1.2 {https://github.com/lh3/seqtk} for extracting reads. The resulting demultiplexed files were processed by the Immcantation pipeline (docker container v4.4.0)^93,94^. The reads were assembled using the AssemblePairs module followed by filtering with the FilterSeq module with -q set to 20. The ParseHeaders module was used to add the sample name to the sequences. Duplicates were removed using the CollapseSeq module with the -–uf set to the 5’ random nucleotide sequence. SplitSeq was used to filter sequences with -DUPCOUNT of 2. IgBLAST v1.21.0^95^ was used to annotate the sequences using the germline database from Cirelli et al 2019^38^. The MakeDB module was used to create ChangeO format files from the IgBLAST output. The functional sequences were selected using ParseDB module to select productive sequences. The D gene usage frequencies were determined using the countGenes function from alakazam v1.3.0 package^94^.

### Activation-induced marker (AIM) and intracellular cytokine staining (ICS) assay to detect antigen-specific CD4^+^ T cells

Antigen-induced marker-based detection of antigen-specific T cells was performed similar to previously described protocols^96,97^. Cryopreserved PBMCs were thawed and washed in RPMI media containing 10% FBS, 1x penicillin/streptomycin and 1x GlutaMAX (R10 media). Cells were counted and then seeded at 1 million cells per well in a round-bottom 96-well plate. Cells were blocked with 0.5 μg/mL anti-CD40 mAb (Miltenyi Biotec) and incubated with anti-CXCR5 and CCR7 for 15 minutes at 37°C. Then, cells were stimulated for 24 hours with one of the following conditions: (1) 5 µg/mL N332-GT5 Env peptide pool “Env”; (2) 1 ng/mL staphylococcal enterotoxin B (SEB) used as a positive control; or (3) DMSO as a negative, unstimulated control plated in duplicate. N332-GT5 Env peptide pools consist of overlapping 15-mer peptides that cover the entire protein sequence and were resuspended in DMSO. An equimolar amount of DMSO is present in both the peptide pool and the unstimulated, negative control. After 24 hours of incubation, intracellular transport inhibitors – 0.25 μL/well of GolgiPlug (BD Biosciences) and 0.25 μL/well of GolgiStop (BD Biosciences) – were added to the samples along with the AIM marker antibodies (CD25, CD40L, CD69, OX40, 4-1BB) and incubated for 4 hours. After, cells were incubated for 15 min at RT with FC Block (Biolegend) and Fixable Live/Dead Blue (Invitrogen). ells were washed with FACS buffer (2% FBS in PBS) and stained with the surface antibodies for 30 minutes at 4°C. Following, cells were fixed with 4% formaldehyde and permeabilized with a saponin-based buffer and subsequently stained with the intracellular cytokine panel for 30 minutes at RT. Finally, the stained cells were washed and acquired on the Cytek Aurora (Cytek Biosciences). Antibody panels and reagents are summarized in table below.

For data analysis, antigen-specific T cells were measured as background subtracted data, where the linear averages of the DMSO background signal, calculated from duplicate wells for each sample, were deducted from the stimulated signal. A minimum threshold for DMSO background signals was set at 0.005% and the limit of quantitation (LOQ) was defined as the geometric mean of all DMSO samples. For each sample, the stimulation index (SI) was calculated as the ratio between the AIM^+^ response in the stimulated condition and the average DMSO response for the same sample. Samples with an SI lower than 2 for CD4 or 3 for CD8 T cells responses and/or with a background subtracted response lower than the LOQ were considered as non-responders. Non-responder samples were set at the baseline, which is the closest log_10_ value lower than the LOQ.

### AIM/ICS Staining Panel

| Antibodies/ Reagents | Clone | Source | Catalog # | Dilution |
| --- | --- | --- | --- | --- |
| LIVE/DEAD Fixable Blue | - | Invitrogen | L23105 | 1/500 |
| GolgiPlug | - | BD Biosciences | 555029 | 1/1000 |
| GolgiStop | - | BD Biosciences | 554724 | 1/1000 |
| Fc Block | - | BioLegend | 422302 | 1/20 |
| Mouse anti-human CD40 | HB14 | Miltenyi | 130-094-133 | 1:200 |
| Mouse anti-human CXCR5 PE-Cy7 | MU5UBEE | Thermo Fisher Scientific | 25-9185-42 | 1:100 |
| Mouse anti-human CCR7 BV650 | G043H7 | BioLegend | 353233 | 1:100 |
| Mouse anti-human CD69 PE-Cy5 | FN50 | BioLegend | 310908 | 1:250 |
| Mouse anti-human CD137 (4-1BB) BV421 | 4B4-1 | BioLegend | 309819 | 1:250 |
| Mouse anti-human CD25 BV605 | BC96 | BioLegend | 302631 | 1:250 |
| Mouse anti-human CD40L BB515 | 24-31 | BD Biosciences | 568170 | 1:250 |
| Mouse anti-human CD134 (OX40) PE | L106 | BD Biosciences | 340420 | 1:250 |
| Mouse anti-human CD8 BUV496 | RPA-T8 | BD Biosciences | 612943 | 1:100 |
| Mouse anti-human CD14 APC-Cy7 | M5E2 | BioLegend | 301820 | 1:100 |
| Mouse anti-human CD16 APC-eFluor780 | eBioCB16 | Thermo Fisher Scientific | 47-0168-42 | 1:100 |
| Mouse anti-human CD20 APC-Cy7 | 2H7 | BioLegend | 302314 | 1:100 |
| Mouse anti-human CD3 BUV395 | SP34-2 | BD Biosciences | 564117 | 1:100 |
| Mouse anti-human CD4 PerCP-Cy5.5 | OKT4 | BioLegend | 317428 | 1:100 |
| Mouse anti-human PD-1 BV785 | EH12.2H7 | BioLegend | 329929 | 1:100 |
| Mouse anti-human CD45RA PE-CF594 | 5H9 | BD Biosciences | 565419 | 1:100 |
| Armenian Hamster anti-ICOS BV480 | C398.4A | BD Biosciences | 566087 | 1:100 |
| Mouse anti-human IFN-γ BV750 | B27 | BD Biosciences | 566357 | 1:100 |
| Rat anti-human IL-2 BUV737 | MQ1-17H12 | BD Biosciences | 612836 | 1:200 |
| Mouse anti-human TNF-α BV711 | MAb11 | BioLegend | 502940 | 1:200 |
| Mouse anti-human Granzyme B Alexa Fluor 700 | GB11 | BD Biosciences | 560213 | 1:1000 |
| Mouse anti-human IL-21 Alexa Fluor 647 | 3A3-N2.1 | BD Biosciences | 560493 | 1:200 |

### PBMC and FNA Flow Cytometry

Frozen PBMC and LN FNA samples were thawed in RPMI media with 10% FBS, 1% GlutaMAX, and 1% Penicillin/Streptomycin (R10). The recovered cells were then stained with the appropriate antibody panel. Fluorescent antigen probes were constructed by combining fluorophore-conjugated streptavidin and biotinylated N332-GT5 (N332-GT5 WT) and N332-GT5 epitope knock-out (N332-GT5 KO) probes in small increments across 45 minutes with an appropriate volume of PBS at room temperature (RT). Thawed cells were incubated with KO probes for twenty minutes at 4°C then incubated with WT probes for thirty minutes at 4°C. A master mix of surface staining antibodies was then added and incubated for thirty minutes at 4°C (see below for PBMC and FNA panels). Cells were prepared for acquisition for either analysis experiments on Cytek Aurora (Cytek Biosciences) or sorting experiments on BD FACSymphony S6 (BD Biosciences). For sorting experiments, 2 μg per 5 million cells of anti-human Total-Seq-C hashtag antibodies were added to each sample following the surface staining master mix during the staining protocol.

For analysis of bulk BGC and GC-TFH a threshold of 250 B cells and 500 CD4+ T cells was used, respectively. For analysis of Env-binding BGC, a threshold of 75 BGC cells was used. LODs for antigen-and epitope-specific Bmem and BGC cells were determined by calculating the mean of 7 pre-immunizations samples.

### PBMC Staining Panel

| Reagent | Clone | Catalog # | Dilution |
| --- | --- | --- | --- |
| AF647 Streptavidin | - | BioLegend (405237) | - |
| BV421 Streptavidin | - | BioLegend (405225) | - |
| PE Streptavidin | - | BioLegend (405245) | - |
| Viability APC-eFluor506 | - | Thermo Fisher Scientific (65-0866-18) | 1:500 |
| CD8a APC-eFluor780 | RPA-T8 | BD Biosciences (563256) | 1:100 |
| CD16 APC-eFluor780 | CB16 | Thermo Fisher Scientific (47-0168-42) | 1:100 |
| CD14 APC-Cy7 | M5E2 | BioLegend (301820) | 1:100 |
| CD3 APC-Cy7 | SP-34 | BD Biosciences (557757) | 1:100 |
| CD20 BUV395 | 2H7 | BD Biosciences (563781) | 1:100 |
| IgG BV605 | G18-145 | BD Biosciences (563246) | 1:100 |
| CD27 PE-Cy7 | O323 | Thermo Fisher Scientific (25-0279-42) | 1:50 |
| IgM PerCP-Cy5.5 | G20-127 | BD Biosciences (561285) | 1:50 |
| IgD AF488 | polyclonal | Southern Biotech (2030-30) | 1:50 |

FNA Staining Panel
| Reagent | Clone | Catalog # | Dilution |
| --- | --- | --- | --- |
| AF647 Streptavidin | - | BioLegend (405237) | - |
| BV421 Streptavidin | - | BioLegend (405225) | - |
| PE Streptavidin | - | BioLegend (405245) | - |
| Viability APC-eFluor506 | - | Thermo Fisher Scientific (65-0866-18) | 1:500 |
| CD4 BV711 | OKT4 | BioLegend (317440) | 1:100 |
| CD8a APC-eFluor780 | RPA-T8 | Thermo Fisher Scientific (47-0088-42) | 1:100 |
| CD16 APC-eFluor780 | CB16 | Thermo Fisher Scientific (47-0168-42) | 1:100 |
| CD20 AF488 | 2H7 | BioLegend (302316) | 1:100 |
| CD38 PE-Cy5 | OKT10 | Produced in house | 1:100 |
| CD3 BUV395 | SP-34 | BD Biosciences (564117) | 1:40 |
| IgG BUV737 | G18-145 | BD Biosciences (612819) | 1:40 |
| IgM PerCP-Cy5.5 | G20-127 | BD Biosciences (561285) | 1:40 |
| PD1 BV605 | EH12.2H7 | BioLegend (329924) | 1:20 |
| CXCR5 PE-Cy7 | Mu5UBEE | Thermo Fisher Scientific (25-9185-42) | 1:20 |
| CD71 PE-CF594 | L01.1 | BD Biosciences (Custom) | 1:20 |

### Single B cell sequencing

After sorting, antigen specific PBMC and LN FNA B cells were wash with PBS and loaded onto a Chromium chip and controller following the manufacturers protocol (10X Genomics). Indexed V(D)J, Feature Barcode, and Gene Expression libraries were prepared following the 10X Genomics protocol with the Feature Barcoding Kit (10X Genomics) and sequenced on a NovaSeq 6000 (Illumina). Demultiplexing and FASTQ generation were done using Cell Ranger v7.2 (10X Genomics). VDJ reads were assembled de novo using Cell Ranger and contigs were aligned to a custom *Macaca mulatta* germline VDJ reference using IgBLAST and the Change-O 1.3.0 package from the Immcantation framework^94^. Sequences were assigned to animals using the MULTIseqDemux function in Seurat v4^98^. Inferred germline sequences (CreateGermline.py) and clones (DefineClones.py) were determined with Change-O. SHM was quantified using the observedMutations command in SHazaM package v1.2.0 comparing HC sequences to the inferred germline sequences (masking the D gene sequences). For clonal abundance and diversity, clonotypes for each animal were quantified using the countClones function in Alakazam package v1.3.0, only animals that had 5 or more clones were included in abundance and diversity calculations. Clonal richness (Chao1) was calculated using the iNEXT package v3.0.0 in R^99^, clonal richness was only calculated for animals that had at least 1 sequence recovered at that timepoint.

### ELISPOT

The antibody-secreting cells (ASCs) in NHP bone marrow were quantified using the enzyme-linked immunosorbent spot (ELISpot) assay as described previously with slight modifications^100^. Briefly, 96-well multi-screen HTS filter plates (MSHAN4B50, Millipore) were coated for 16 hours at 4°C with either 5μg/mL of Goat Anti-Human Ig-UNLB (Southern Biotech, Cat# 2010-01) for capturing total rhesus IgG or 20μg/mL of Galanthus Nivalis Lectin (Vector laboratories, Cat# L-1240) for capturing N332 antigen. The plates were then washed four times with sterile PBS and blocked with R10 media for 2 hours in a 5% CO2 incubator at 37°C. For antigen capture, the GNL-coated plate was further incubated with 10μg/mL of N332 antigen for 2 hours at 37°C. Bone marrow (BM) mononuclear cells were harvested using Ficoll Paque Plus from 2 mL of BM aspirates as reported (Kasturi SP et al 2020, Sci Immunol). 100μl of cells (5×106 cells/mL) were added to the wells in the first row in duplicates (~333K cells), and serially diluted three-fold, and incubated overnight for ~16 hours in a 5% CO2 incubator at 37°C. Following incubation, the plates were washed four times with PBS followed by four washes with PBS containing 0.05% Tween 20 (PBS-T). Goat Anti-Human IgG-biotin (Southern Biotech, Cat# 2045-08) was added at a dilution of 1:500 and incubated for two hours at room temperature. Plates were washed with PBS followed by PBS-T four times each and Avidin D-HRP (Vector Labs) was added at a dilution of 1:1000 and incubated for 1 hour at room temperature. Finally, plates were washed four times in PBS-T followed by four washes with PBS, and spots were developed for 5 minutes using a filtered 3-amino 9-ethylcarbazole (AEC) substrate (0.3 mg/mL AEC in 0.1 M sodium acetate buffer, pH 5.0, with a 1:1000 dilution of 3% hydrogen peroxide). The reaction was stopped by washing the plates with running tap water, and air-dried plates before imaging using the Immunospot CTL counter and Image Acquisition 4.5 software (Cellular Technology). Total IgG+ ASC or Env-specific ASCs/spots were counted manually using magnified images and reported as the absolute number of ASCs per million BM cells or % of antigen specific ASCs as a fraction of total ASCs.

### Antibody production

High-throughput antibody production was performed as previously described^61,101^. Briefly, antibodies were produced using the ExpiCHO Expression System (Thermo Fisher) in baffled 96 deep-well blocks (Thomas Scientific) according to manufacturer’s instructions, with the following adjustments: the cell density was increased to 10^7^ viable cells/mL, cell culture volumes were increased to 1 mL/well, ExpiFectamine Reagent (Thermo Fisher) volume was increased to 6 μL/well, and incubator shake speed was set to 1300 RPM. After 7 days incubation, supernatants were supplemented with 40 μL 1M Tris, pH 8.0, harvested by centrifugation and purified using Protein A HP MultiTrap plates (GEHealthcare) per manufacturer’s instructions.

### Antibody binding affinities

We measured kinetics and affinity of antibody-antigen interactions on Carterra LSA using CMDP Sensor Chip (Carterra). This chip type has lower ligand capacity and excellent diffusion characteristics. It is recommended for use when reducing avidity is necessary. 1x HBS-EP+ pH 7.4 was our running buffer (20x stock from Teknova, Cat. No H8022) supplemented with BSA at 1mg/ml. We followed Carterra software instructions to prepare chip surface for ligands capture. In a typical experiment about 500-700 RU of capture antibody (SouthernBiothech Cat no 2047-01) in 10 mM Sodium Acetate pH 4.5 was amine coupled. The critical detail here was the concentration range of the amine coupling reagents and capture antibody. We used N-Hydroxysuccinimide (NHS) and 1-Ethyl-3-(3-dimethylaminopropyl) carbodiimidehydrochloride (EDC) from Amine Coupling Kit (GE order code BR-1000-50). As per kit instruction 22-0510-62 AG the NHS and EDC should be reconstituted in 10 ml of water each to give 11.5 mg/ml and 75 mg/ml respectively. However, the highest coupling levels of capture antibody were achieved by using 10 times diluted NHS and EDC during surface preparation runs. Thus, in our runs the concentrations of NHS and EDC were 1.15 mg/ml and 7.5 mg/ml. The activation time was reduced to 1min. The concentrated stocks of NHS and EDC could be stored frozen in −20C for up to 2 months without noticeable loss of activity. The SouthernBiotech capture antibody was used at concentration 25ug/ml with 10 minutes contact time. Phosphoric Acid 1.7% was our regeneration solution with 60 seconds contact time and injected three times per each cycle. Solution concentration of ligands was around 1 ug/ml and contact time was 5 min. Raw sensograms were analyzed using Kinetics software (Carterra), interspot and blank double referencing, Langmuir model. Analyte concentrations were quantified on NanoDrop 2000c Spectrophotometer using Absorption signal at 280 nm. For best results analyte samples should be buffer exchanged into the running buffer using dialysis. We typically cover broad range of affinities in our runs and the best referencing practices are different depending on how fast the off-rate is for particular ligand. For fast off-rate (faster than 9e-3 1/s) we use automated batch referencing that includes overlay y-aline and higher analyte concentrations. For slow off-rates (9e-3 1/s or less) we use manual process referencing that includes serial y-align and lower analyte concentrations. After automated data analysis by Kinetics software, we also did additional filtering to remove datasets with highest response signals smaller than signals from negative controls. This additional filtering could be performed automatically using R-script.

### Statistical Analysis

Statistical tests were carried out using GraphPad Prism 10. For most graphs unpaired two-tailed Mann-Whitney tests were used for the following comparisons (i) G1 vs 2, G1 vs 3, G1 vs 4, G1 vs 5, G1 vs 6, (ii) G2 vs 5, G2 vs 6, (iii) G4 vs 5, (iv) G5 vs 6. Only (i) tests were done for the BG18 sequencing frequency graphs except in Figure 3A were done using Fishers exact tests comparing each group to G1 individually. P-values are listed if <0.1 or reported as NS if >0.1. Additional tests were carried out in Figure 4 comparing each group to the pre-immunization samples using a Kruskal-Wallis test without correction. Those p-values are represented by the following scale: NS > 0.05, * <0.05, ** <0.01, *** <0.001, **** <0.0001. All p-values reported in Figure 7 are from Mann-Whitney tests.

## Material availability

This study did not generate new unique reagents

## Data and code availability

- All sequencing files will be uploaded to SRA
- All original code will be publicly available on Github
- Any additional information required to reanalyze the data reported in this paper is available from the lead author upon request

## Acknowledgments

We thank Jennifer S Wood and the staff of the Emory National Primate Research Center. We are grateful to the Flow Cytometry and Next Generation Sequencing core facilities at La Jolla Institute for Immunology for their services.

## Funding

This work was funded by the National Institute of Allergy and Infectious Diseases of the NIH (NIAID-NIH) under award CHAVD UM1AI144462. This research was funded in part by the Emory National Primate Research Center Grant Nos. ORIP/OD P51OD011132 and U42 PDP11023. The Emory National Primate Center is supported by the National Institutes of Health, Office of Research Infrastructure Programs/OD [P51OD011132 and U42 PDP11023]. The Emory Primate Genomics Core received funding from P51OD011132 and S10OD026799. The NovaSeq 6000 used in this work was funded by NIH grant S10OD025052.

## Author Contributions

Conceptualization, P.J.M., D.J.I., and S.C.; Methodology, P.J.M., E.M.Z., K.A.R., A.A.U., S.S., A.P., S.P.K., C.T.W., S.E.B., D.J.I., and S.C.; Investigation, P.J.M., E.M.Z., K.A.R., J.M.S., M.S., K.N., K.K.M., L.M., A.A.U., S.S., A.P., O.K., A.L., P.G.L., I.P., A.M., O.L.R., K.S., S.S., M.L.S., E.B.A., W.P., J.G., and S.X.; Resources, K.A.R., J.M.S., K.N., N.P., E.G., N.A., M.K., D.L., S.E., B.S.H., D.L., V.R.L., E.B.A., W.P., J.G., S.X., D.C.C., S.P.K., C.T.W., S.E.B., G.S., W.R.S., D.J.I., and S.C.; Writing-Original Draft, P.J.M., D.J.I., and S.C.; Writing-Review & Editing, P.J.M., J.M.S., C.T.W., W.R.S., D.J.I., and S.C.; Supervision, S.P.K., C.T.W., S.E.B., G.S., W.R.S., D.J.I., and S.C.; Project Administration, P.J.M., L.M., D.C.C., D.J.I., S.C.; Funding Acquisition, C.T.W., S.E.B., G.S., W.R.S., D.J.I., and S.C..

## Declaration of interests

D.J.I. and S.C. are inventors on a patent for the SMNP adjuvant (US11547672B2). K.A.R. and D.J.I. are inventors on patent applications for the synergistic combination of alum and SMNP adjuvants (PCT/US2022/074302 and US No. 17/816,045). D.J.I. and W.R.S. are named as inventors on a patent for pSer technology (US No. 11,224,648 B2). J.M.S. and W.R.S. are inventors on patent applications related to immunogens in this manuscript that have been filed by Scripps and IAVI. W.R.S. is an employee and shareholder of Moderna, Inc. All other authors declare no competing interests.

## SUPPLEMENTARY FIGURE LEGENDS

**Supplementary Figure 1:**
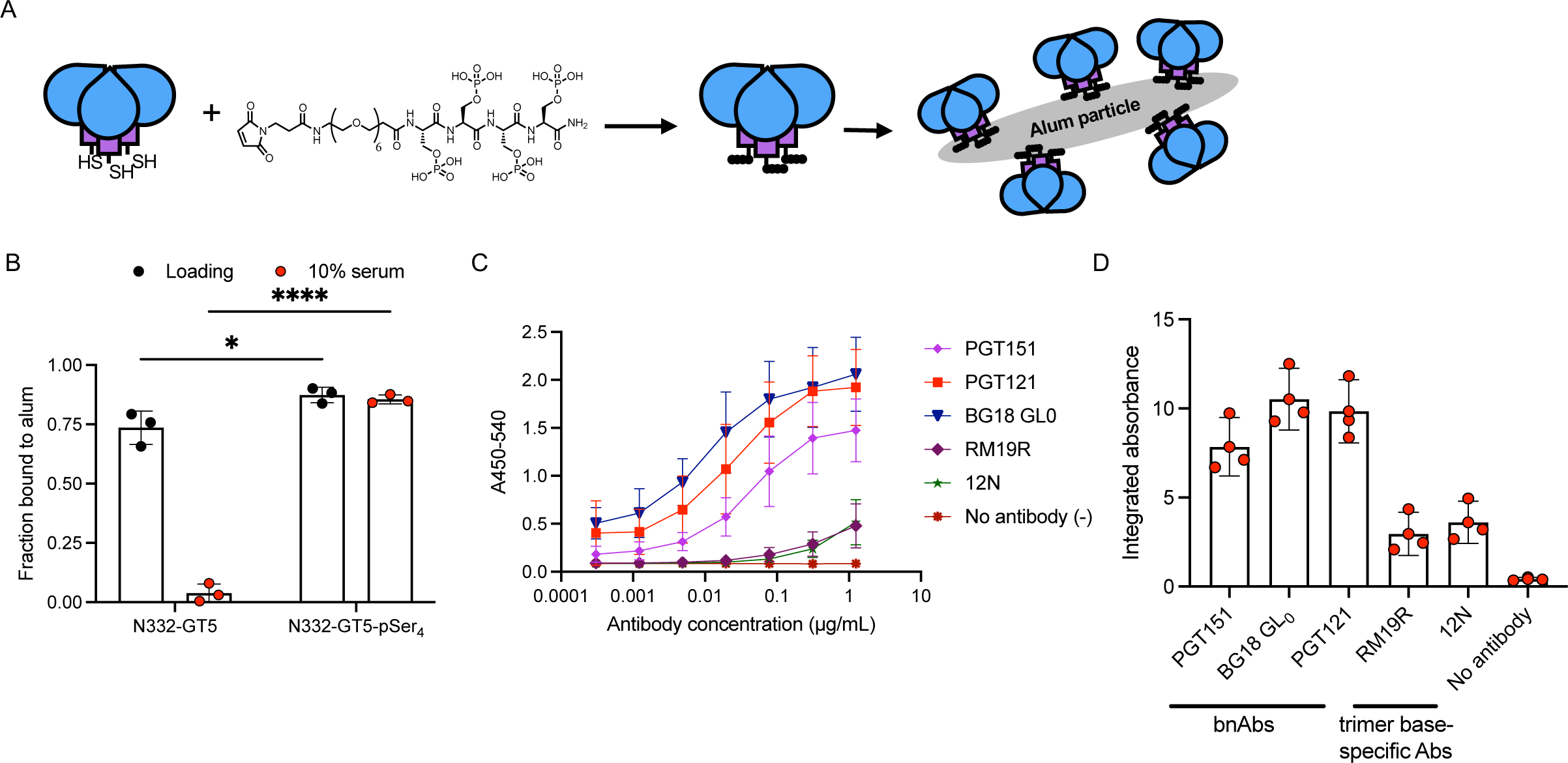
pSer modification and antigenicity. **A)** Schematic of pSer peptide coupling to c-terminal Cys at the base of Env trimers and subsequent binding to aluminum hydroxide particles. **B)** Fluorescently-labeled pSer-conjugated or unmodified N332-GT5 trimers were mixed with alum in TBS, and the fraction of protein bound to alum was assessed after initial 30-minute adsorption (“Loading”) or after 24-hour incubation in 10% mouse serum at 37°C by fluorescence spectroscopy. **C)** Antigenicity profiling ELISA assessing binding of serial dilutions of the indicated mAbs to pSer-N332-GT5 adsorbed to plate-immobilized alum. **D)** Area-under-the-curve values for mAb binding vs. antibody concentration from data shown in (**C**). Mean and SD are plotted. Statistical significance was tested using two-way ANOVA with Sidak’s multiple comparisons test in (**B**). * p < 0.05, ****p < 0.0001.

**Supplementary Figure 2:**
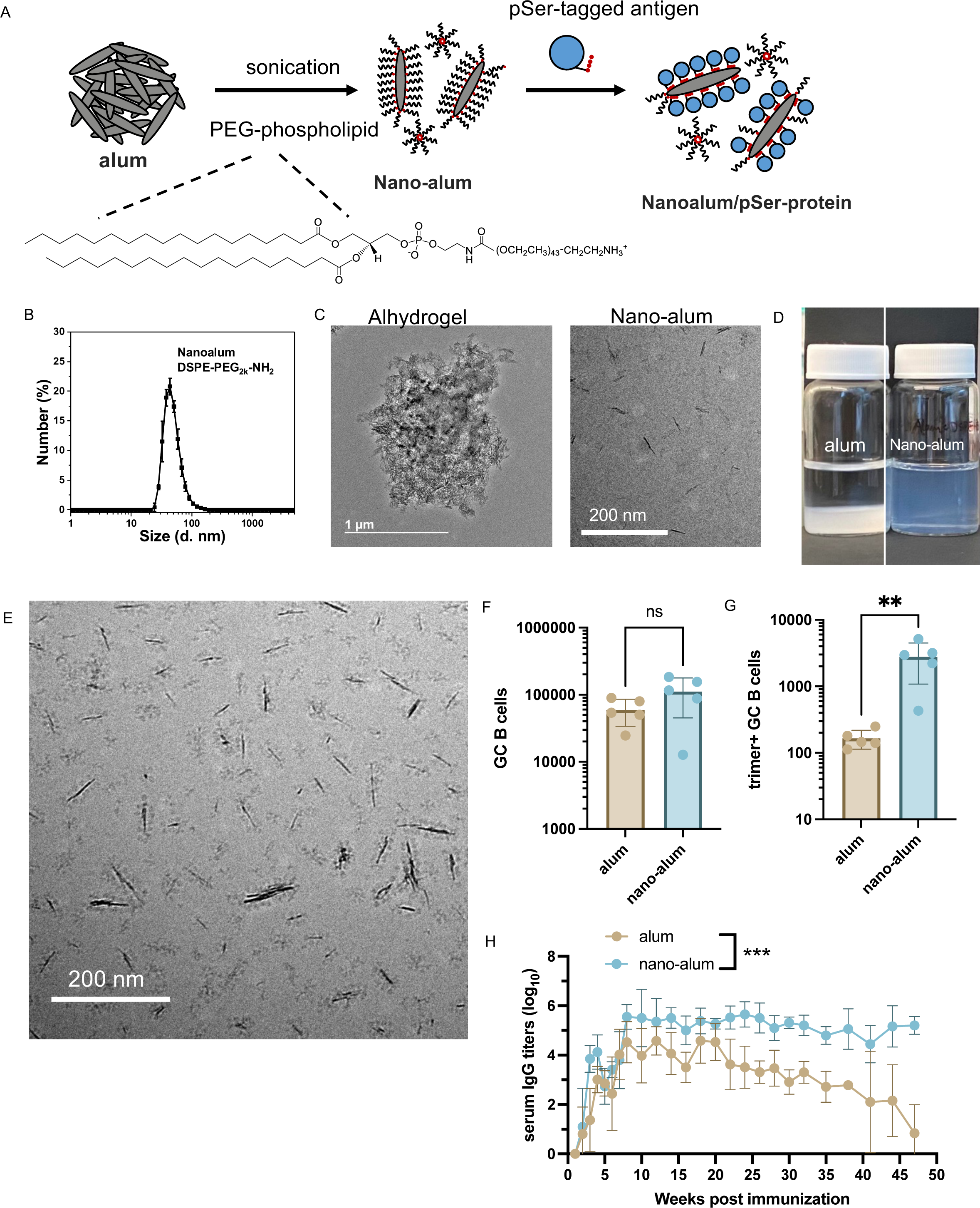
Nano-alum synthesis and characterization. **A)** Schematic of nano-alum formulation and binding with pSer-trimers. **B)** Number-average particle size of nano-alum determined by dynamic light scattering. C) TEM image of neat Alhydrogel and cryoEM image of nano-alum. **D)** Photographs illustrating physical appearance of sedimented alum solution vs. stable opalescent nano-alum solution. **E)** CryoEM image of pSer-trimer mixed with nano-alum showing trimers decorating individual aluminum hydroxide nanocrystals. F-H) BALB/c mice (*n* = 5 animals/group) were immunized with 5 µg pSer-trimer adsorbed to 50 µg Alhydrogel (alum) or 50 µg nano-alum. Shown are flow cytometry analyses of total GC B cells **(F)** and trimer-specific GC B cells **(G)** on day 14 and serum IgG titers determined by ELISA over time (H). Statistical significance determined by two-tailed *t* test in **(F, G)** and two-way ANOVA in (H). ns, not significant; **, p < 0.01; ***, p < 0.001.

**Supplementary Figure 3:**
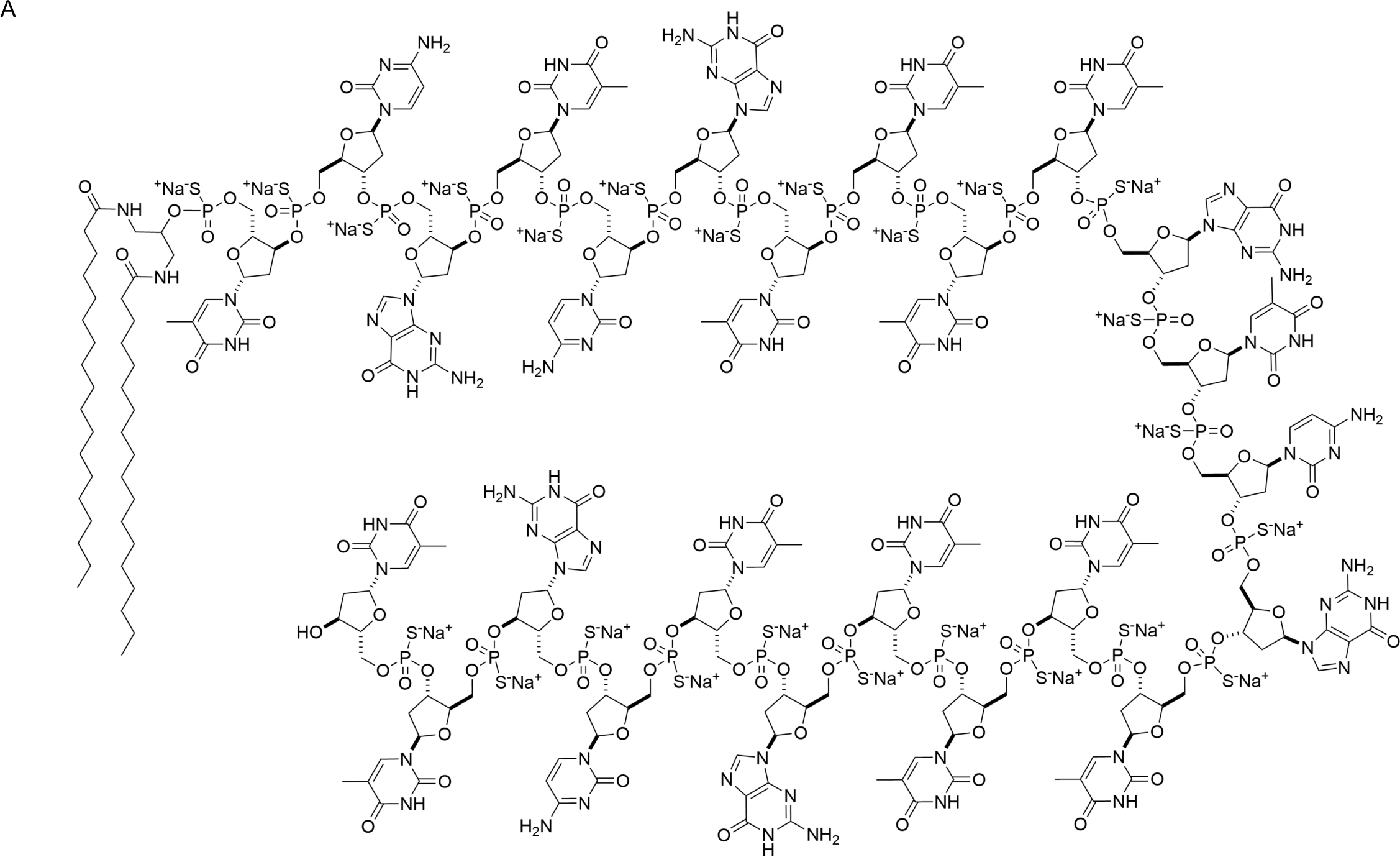
Amph-CpG adjuvant. **A)** Molecular structure of Amph-CpG.

**Supplementary Figure 4:**
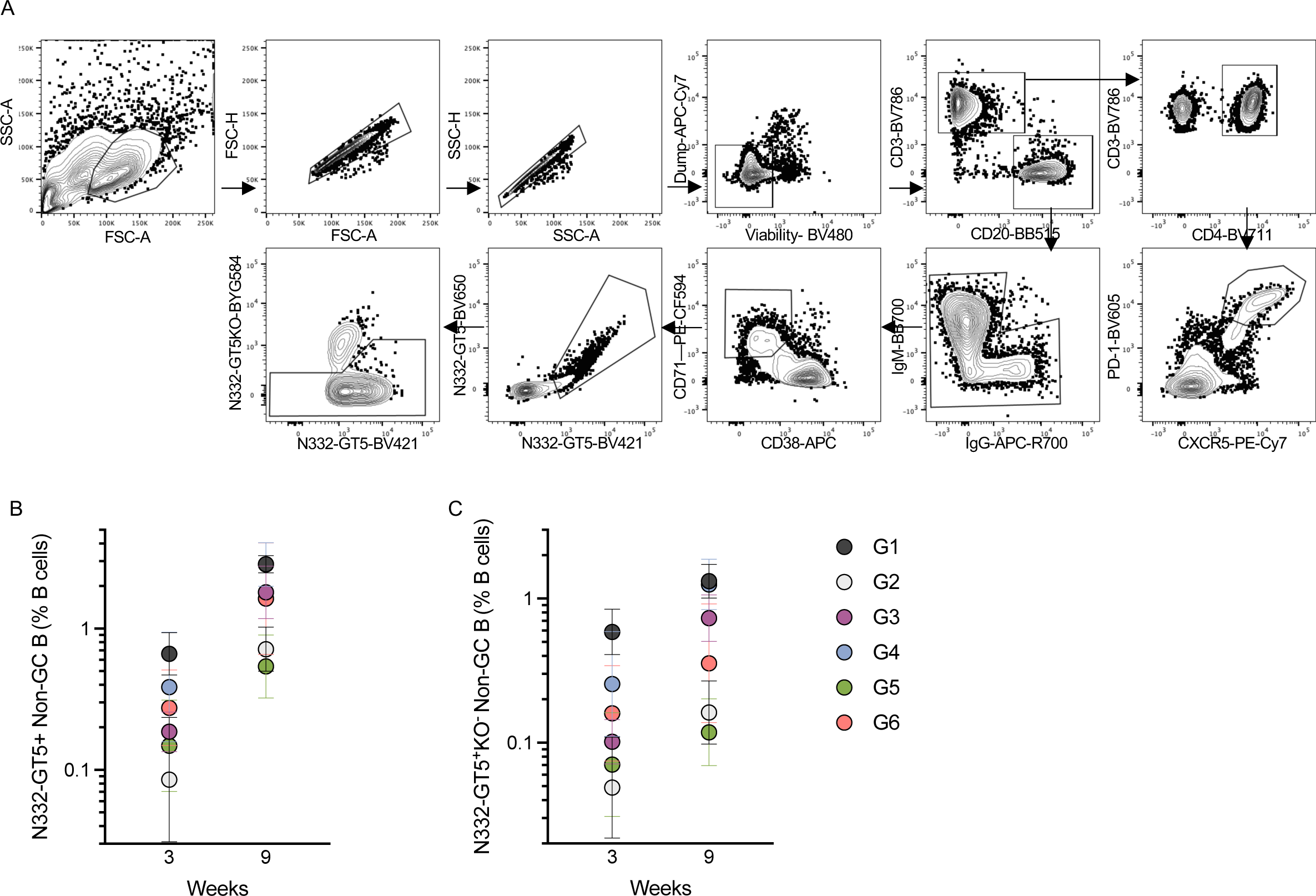
Germinal center flow cytometry gating. **A)** Gating strategy for LN FNA analysis. GC-T_FH_ gating and B_GC_ gates used for quantification are shown with antigen- and epitope-specific gates. Epitope-specific B_GC_ cells were sorted. **B)** Antigen-specific non-GC B cell frequency (N332-GT5-AF647^+^N332-GT5-BV421^+^) as a percentage of total B cells (CD20^+^). Geometric mean and SD are shown for each group at week 3 and 9. **C)** Antigen-specific non-GC B cell frequency (N332-GT5-AF647^+^N332-GT5-BV421^+^N332-GT5KO-PE^−^) as a percentage of total B cells (CD20^+^). Geometric mean and SD are shown for each group at week 3 and 9.

**Supplementary Figure 5:**
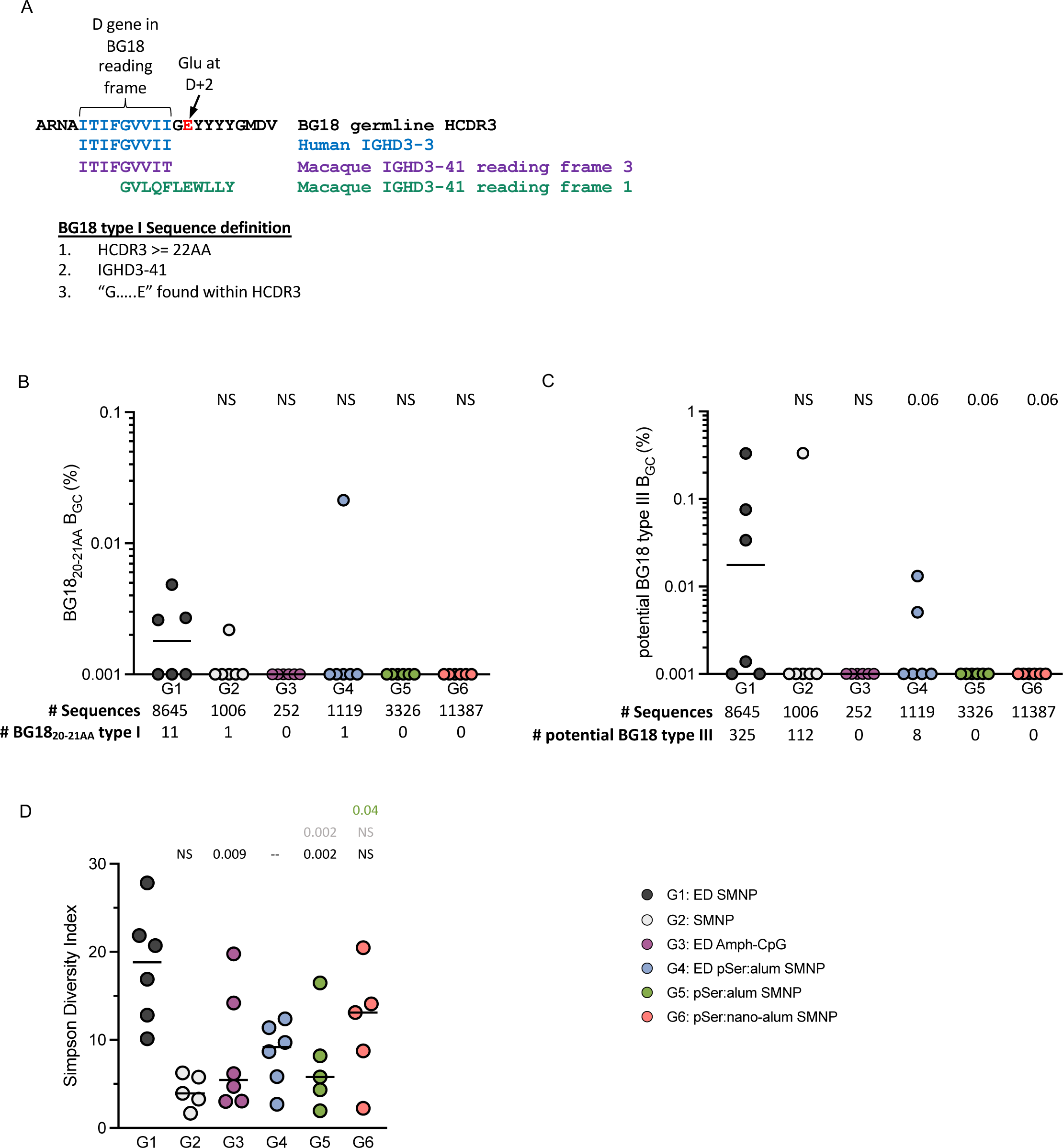
BG18 sequencing. **A)** Sequence alignment of germline BG18 H-CDR3 with macaque IGHD3-41 and sequence definition of BG18 type I BCRs. **B)** Frequency of BG18_short_ B_GC_ cells among total B cells, plotted per animal. Numbers below indicate total number of paired BCR sequences recovered from each group and the total number of BG18_short_ BCRs recovered. **C)** Frequency of potential BG18 type III B_GC_ cells among total B cells, plotted per animal. Numbers below indicate total number of paired BCR sequences recovered from each group and the total number of potential BG18 type III BCRs recovered. **D)** Simpson diversity index of BCR sequences plotted per animal.

**Supplementary Figure 6:**
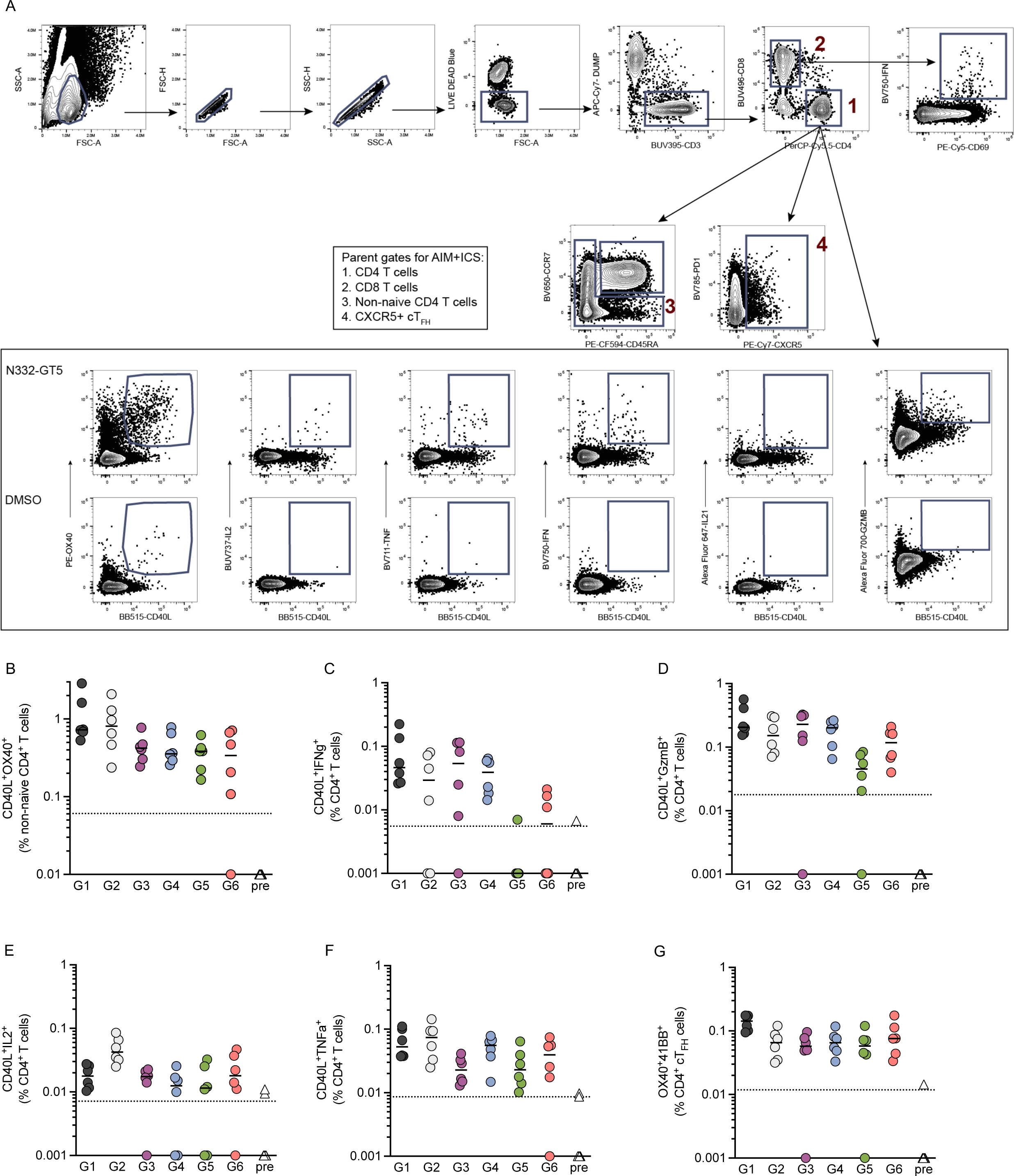
AIM ICS gating and data. **A)** Flow cytometry gating for AIM ICS T cell assay. **B)** Frequency of AIM^+^ (CD40L^+^OX40^+^) T cells out of total CD4^+^ T cells. **C)** Frequency of AIM^+^ICS^+^ (CD40L^+^IFNγ^+^) T cells out of total CD4^+^ T cells. **D)** Frequency of AIM^+^ICS^+^ (CD40L^+^GzmB^+^) T cells out of total CD4^+^ T cells. **E)** Frequency of AIM^+^ICS^+^ (CD40L^+^IL2^+^) T cells out of total CD4^+^ T cells. **F)** Frequency of AIM^+^ICS^+^ (CD40L^+^TNFα^+^) T cells out of total CD4^+^ T cells. **G)** Frequency of AIM^+^ (OX40^+^41BB^+^) cT_FH_ cells out of total CD4^+^ cT_FH_ cells.

**Supplementary Figure 7:**
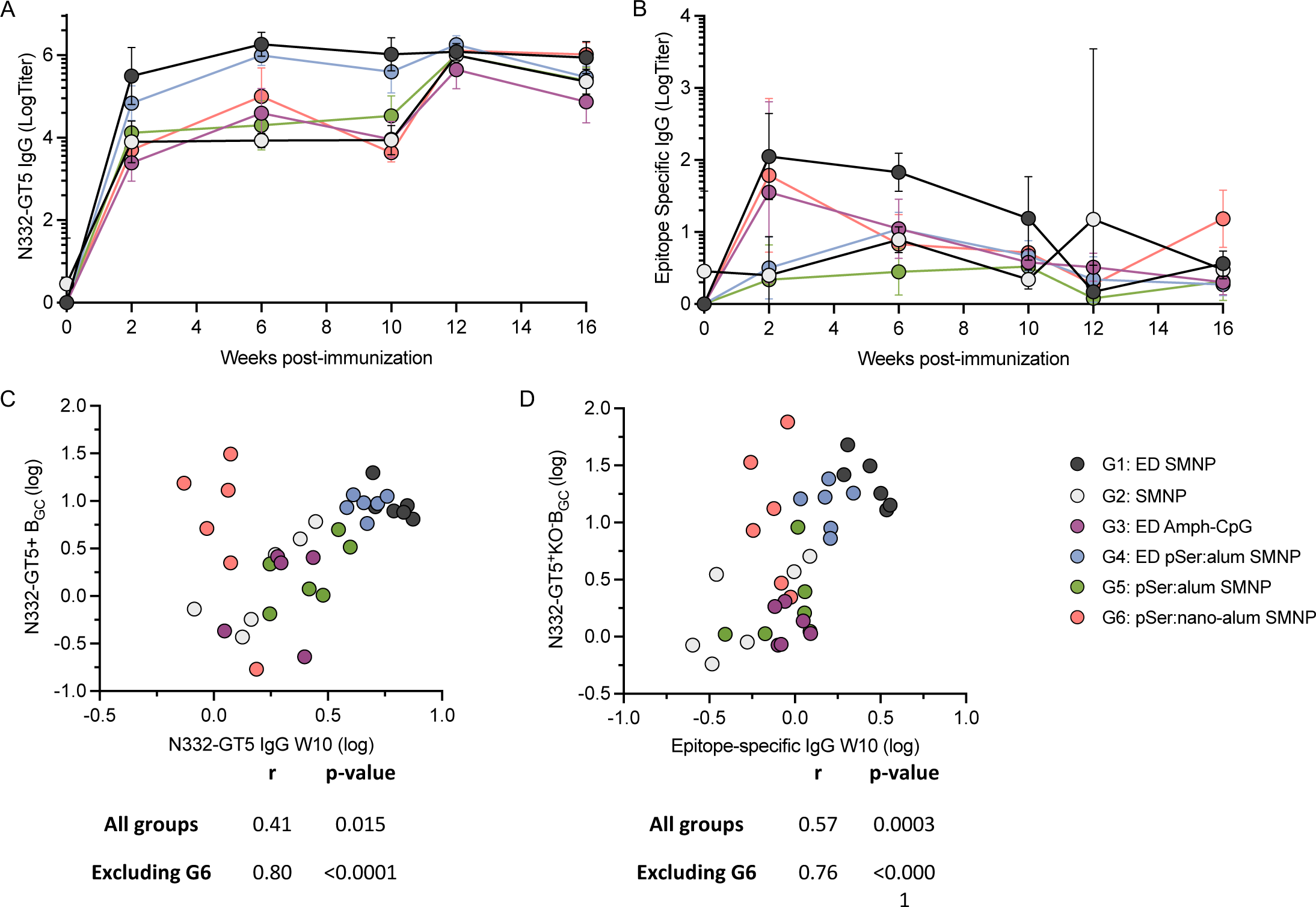
ELISA data and B_GC_ correlations. **A)** Longitudinal endpoint titer curves of total antigen-specific serum IgG measured by ELISA. **B)** Longitudinal endpoint titer curves of total epitope-specific serum IgG measured by ELISA. Geometric mean and SD are plotted for each group at each time point on a Log(titer) axis. **C)** Correlation between total post-prime antigen-specific B_GC_ and week 10 antigen-specific IgG AUC. r and p-value are from pearson correlation analysis run with all groups included or excluding G6. **D)** Correlation between total post-prime epitope-specific B_GC_ and week 10 epitope-specific IgG AUC. r and p-value are from pearson correlation analysis run with all groups included or excluding G6.

**Supplementary Figure 8:**
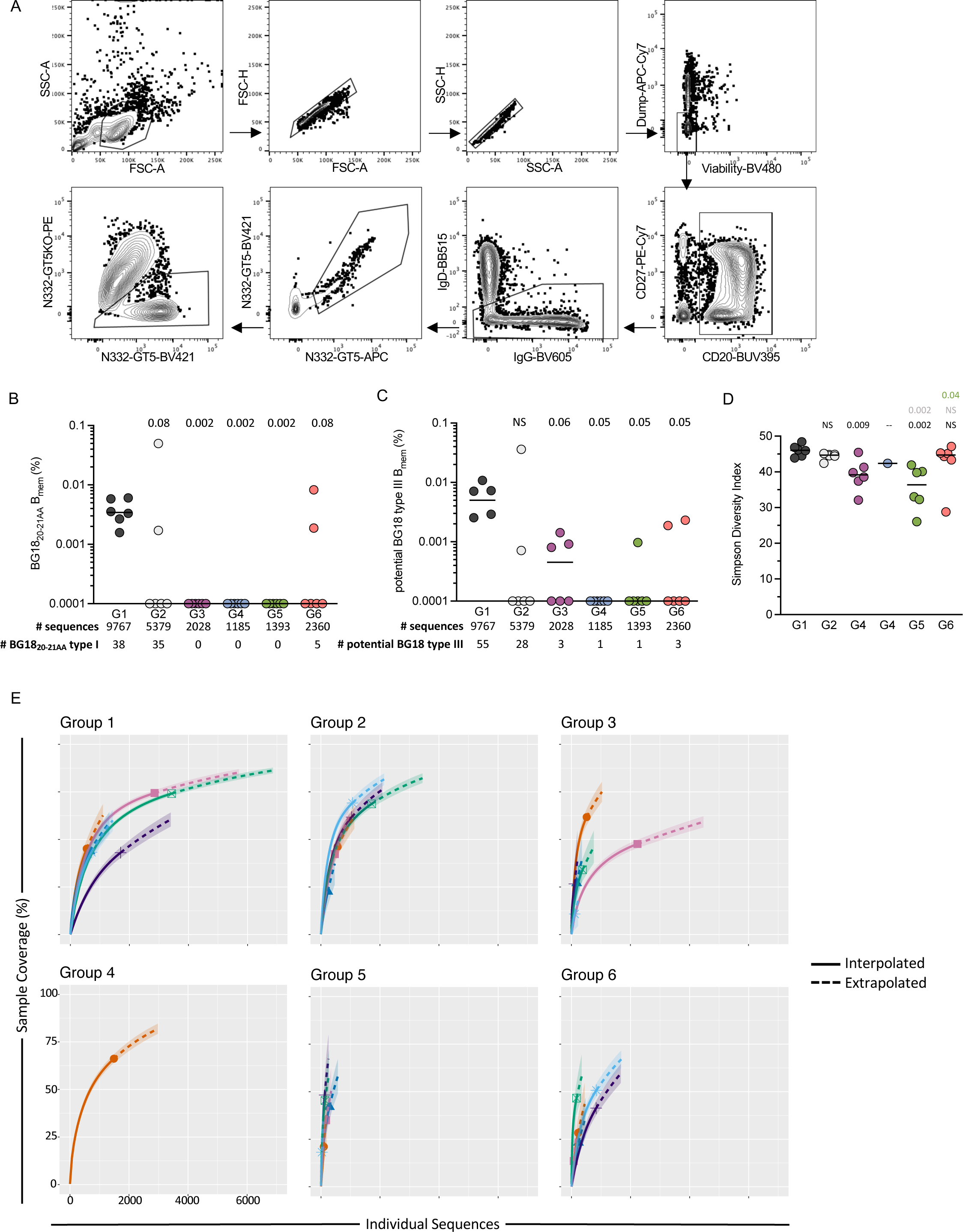
B_mem_ flow cytometry and sequencing. **A)** Flow cytometry gating strategy for isolating epitope-specific Bmem cells from PBMCs. **B)** Frequency of BG18_short_ B_GC_ cells among total B cells, plotted per animal. Numbers below indicate total number of paired BCR sequences recovered from each group and the total number of BG18_short_ BCRs recovered. **C)** Frequency of potential BG18 type III B_GC_ cells among total B cells, plotted per animal. Numbers below indicate total number of paired BCR sequences recovered from each group and the total number of potential BG18 type III BCRs recovered. **D)** Simpson diversity index of BCR sequences plotted per animal. Statistical significance was tested using unpaired two-tailed Mann-Whitney tests with the p-values listed on each graph representative of the tests carried out, NS listed when p-value was >0.1. **E)** iNext plots showing the results of rarefaction analysis performed on the week 12 Bmem sequences to determine the % sequence coverage based on the number and clonality of the recovered sequences. Each group is plotted separately with each line representing a single animal. Only animals that had more than 5 unique clones were included in the analysis.

**Supplementary Figure 9:**
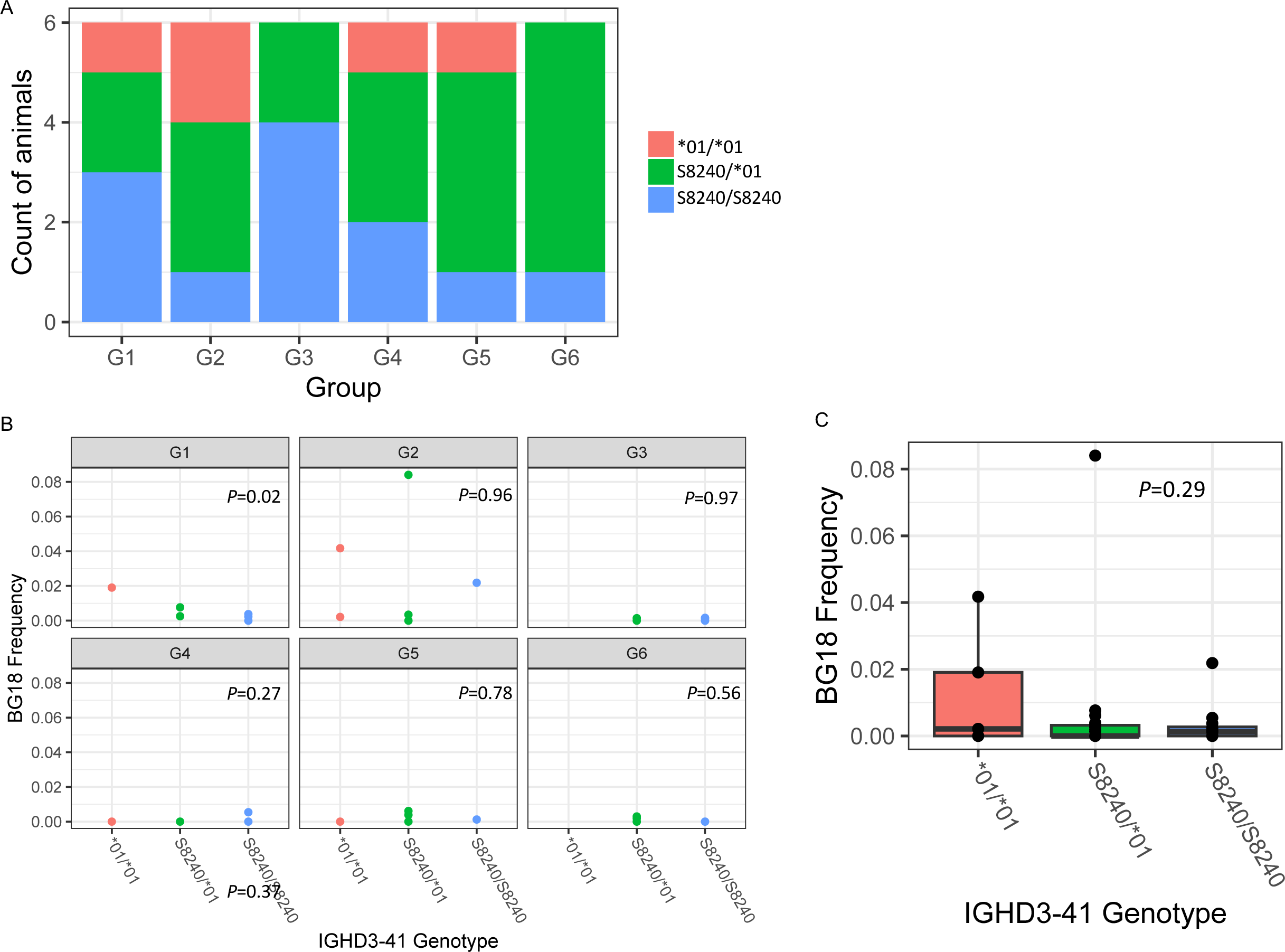
IGHD3-41 Immunogenetics. **A)** IGHD3-41 genotype distribution for each animal separated by group. **B)** Week 12 BG18 type I B_mem_ frequency plotted by genotype and group. Each dot represents an individual animal. **C)** Week 12 BG18 type I B_mem_ frequency plotted by genotype for all 42 animals together. Statistical significance was tested using ANOVA with the p-values listed on each graph. For B) p-values represent within-group comparisons.

**Supplementary Figure 10:**
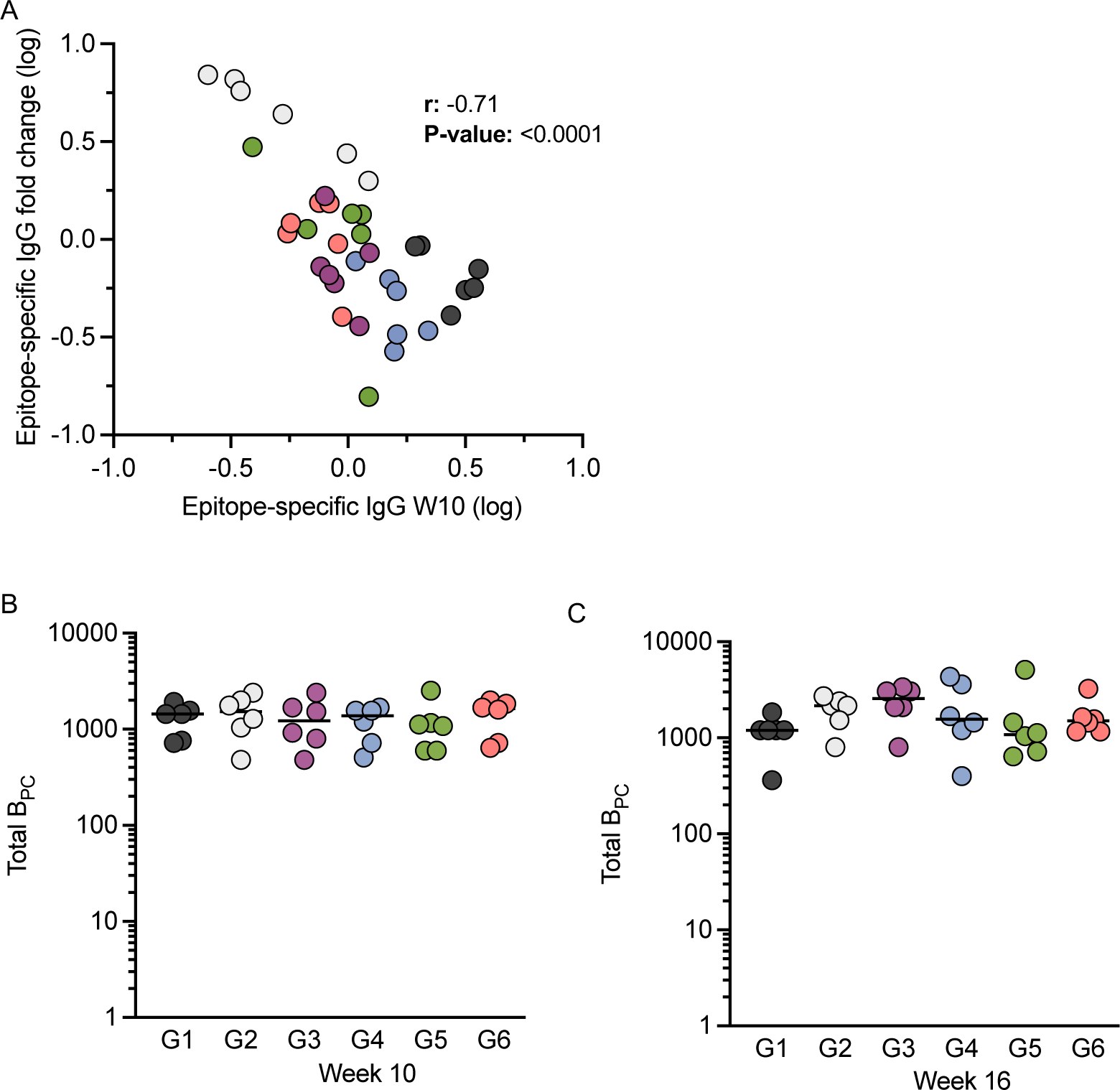
Serum IgG correlations and B_PC_. **A)** Correlation between epitope-specific fold change increase from week 10 to 12 and epitope-specific week 10 AUC. r and p-value are from pearson correlation analysis. **B-C)** Total IgG^+^ BM-B_PC_ measured from bone marrow aspirates by ELISpot assay at weeks 10 and 16.

**Supplementary Figure 11:**
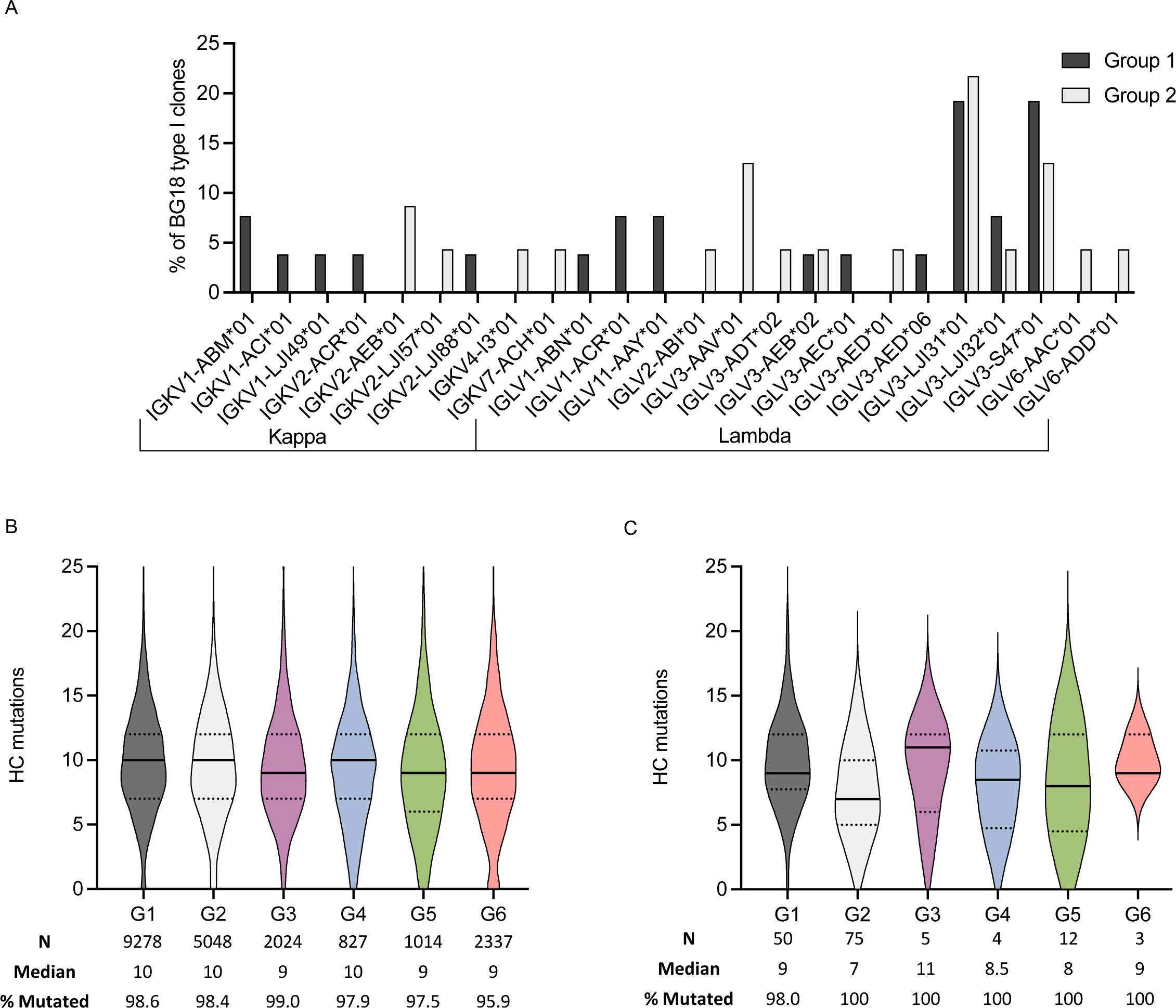
Light chain usage and SHM across groups. **A)** Graph of the distribution of IGHK and IGHL genes used by BG18 type I clones in groups 1 and 2. **B)** Heavy chain nucleotide mutations plotted for each group of total week 12 B_mem_ cells. Number of sequences, median mutation count, and percentage of all cells with 1 or more mutations are listed below graphs. **C)** Heavy chain nucleotide mutations plotted for each group of week 12 BG18 type I B_mem_. Number of sequences, median mutation count, and percentage of all cells with 1 or more mutations are listed below graphs

**Supplementary Table 1.**
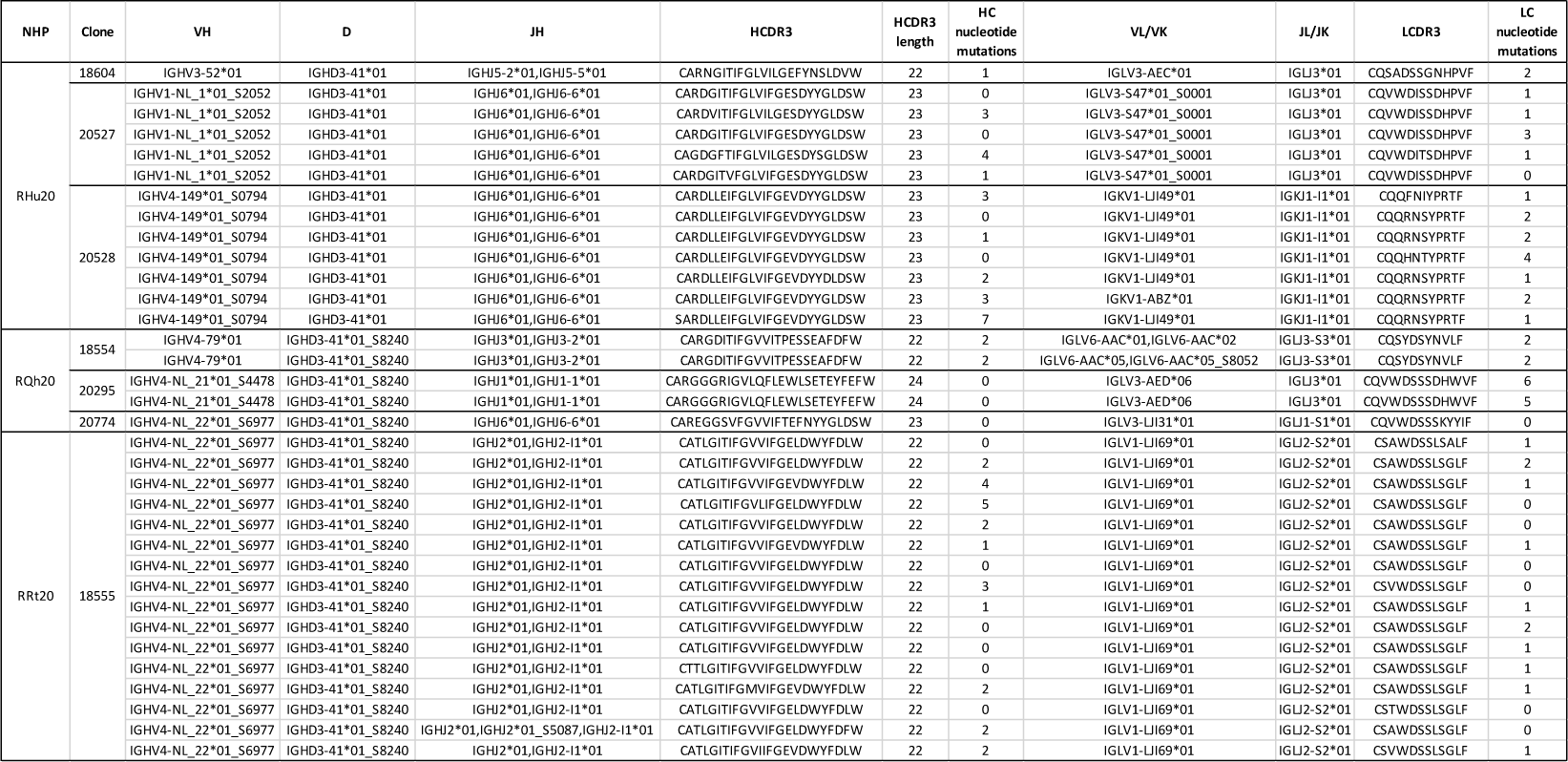
Week 3 BG18 type I B_GC_ sequences.

**Supplementary Table 2.**
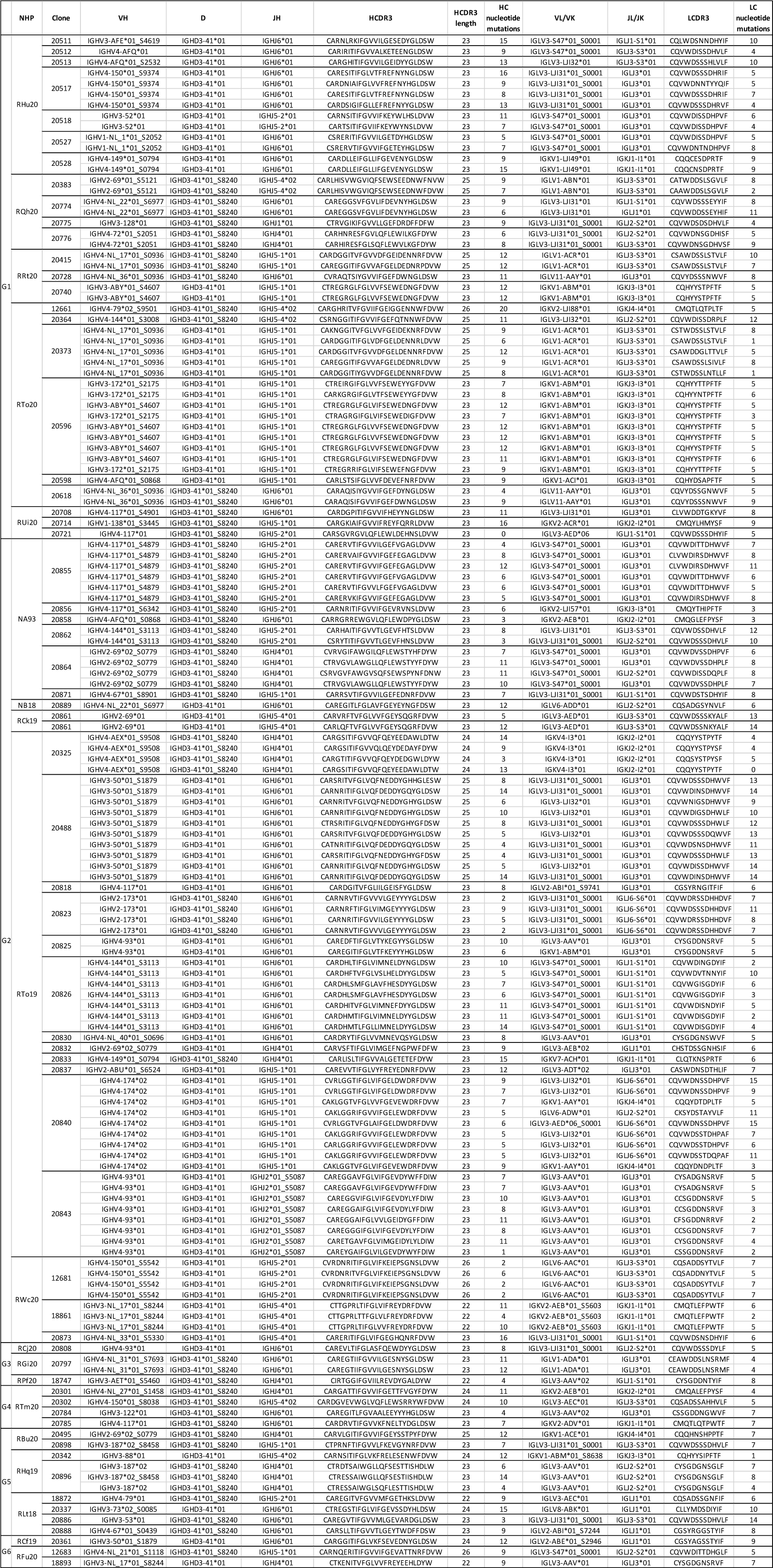
Week 12 BG18 type I B_mem_ sequences.

**Supplementary Table 3.**
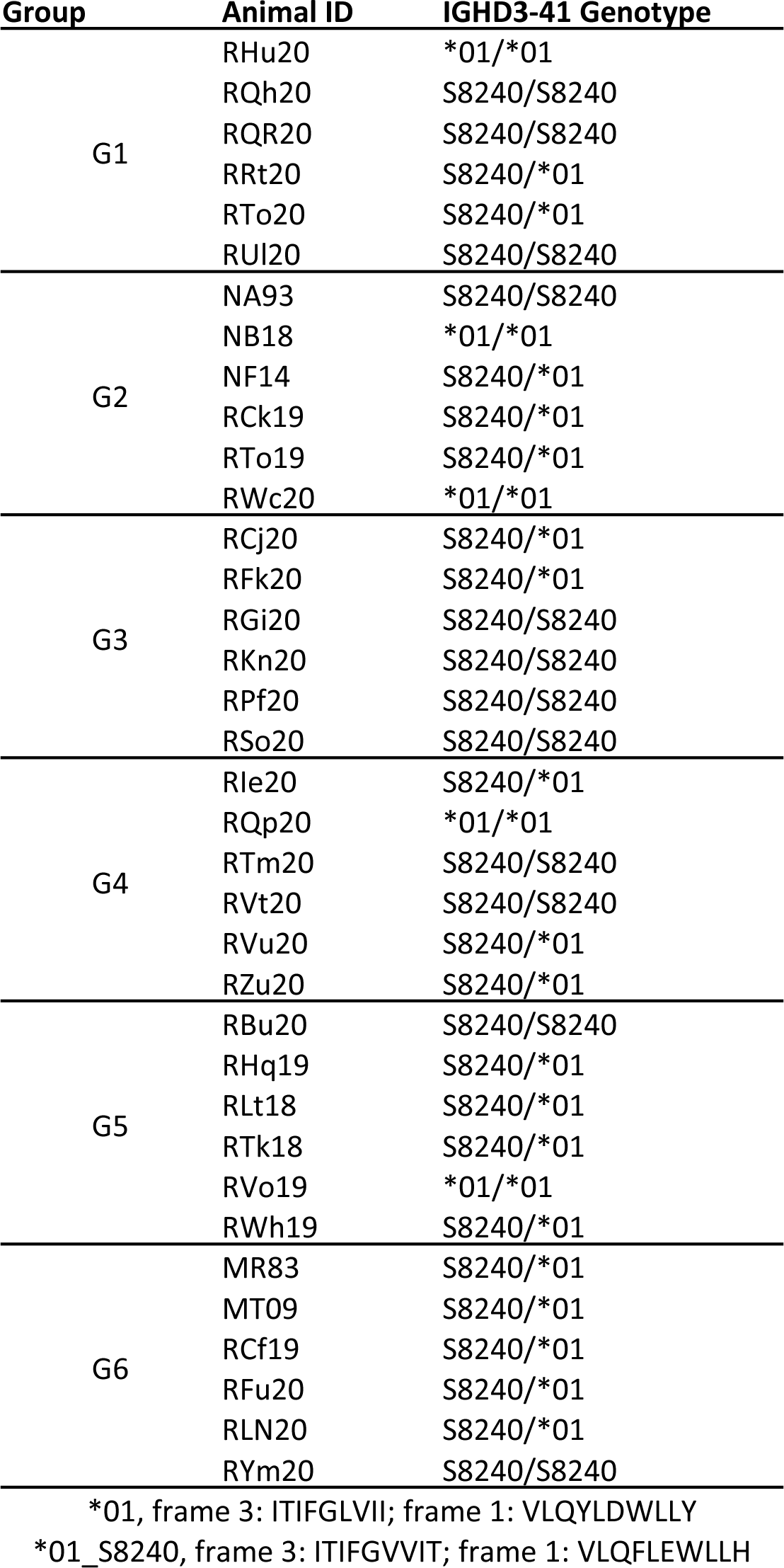
*IGHD3-41* genotypes.

## Notes

### Summary of Updates

Two author names were spelled incorrectly in the original version and have been corrected

